# Genetic reduction of PERK-eIF2α signaling in dopaminergic neurons drives cognitive and age-dependent motor dysfunction

**DOI:** 10.1101/2020.04.07.028241

**Authors:** Francesco Longo, Maria Mancini, Pierre L. Ibraheem, Sameer Aryal, Caterina Mesini, Jyoti C. Patel, Elena Penhos, Nazia Rahman, Maggie Donohue, Emanuela Santini, Margaret E. Rice, Eric Klann

## Abstract

An array of phenotypes in animal models of neurodegenerative disease have been shown to be reversed by neuronal inhibition of PERK, an eIF2α kinase that modulates the unfolded protein response (UPR). This suggests that targeting PERK therapeutically could be beneficial for treatment of human disease. Herein, using multiple genetic approaches we show that selective deletion of the PERK in mouse midbrain dopaminergic (DA) neurons results in multiple cognitive and age-dependent motor phenotypes. Conditional expression of phospho-mutant eIF2α in DA neurons recapitulated the phenotypes caused by deletion of PERK, consistent with a causal role of decreased eIF2α phosphorylation. In addition, deletion of PERK in DA neurons resulted in altered *de novo* translation, as well as age-dependent changes in axonal DA release and uptake in the striatum that mirror the pattern of motor changes observed. Taken together, our findings show that proper regulation of PERK-eIF2α signaling in DA neurons is required for normal cognitive and motor function across lifespan, and also highlight the need for caution in the proposed use of sustained PERK inhibition in neurons as a therapeutic strategy in the treatment of neurodegenerative disorders.

## Introduction

Altered proteostasis and the aggregation of misfolded proteins represent a shared pathological trait across many neurodegenerative diseases^1^. The sustained and inappropriate accumulation of damaged proteins constitutes a pathological load for the endoplasmic reticulum (ER), which results in ER-stress^2^ and activation of the unfolded protein response (UPR), an integrated signaling pathway that protects cells by restoring normal proteostasis. UPR signaling controls the overall burden of misfolded proteins through general translational arrest and increased translation of transcriptional genes that enhance ER protein-folding capacity and quality control through the degradation of proteins with aberrant conformation. Multiple pieces of evidence suggest that the UPR plays a key role in maintaining neuronal function at the level of synapses, connectivity, and brain development^3^.

A critical component of the UPR is protein kinase R-like endoplasmic reticulum kinase (PERK), which phosphorylates the eukaryotic initiation factor 2a (eIF2α) at serine 51, thereby controlling the initiation step of protein synthesis and subsequently preventing an overload of proteins in the ER lumen^4^. Paradoxically, eIF2α phosphorylation via PERK increases the synthesis of transcription factors that contain unread open reading frames (uORFs) in the 5’UTR of their mRNAs, including activating transcription factor 4 (ATF4), which is involved in the expression of several UPR target genes^5^. However, when ER stress is sustained and the adaptive mechanisms of the UPR are not sufficient to recover cellular protein homeostasis, a switch to pro-apoptotic signals triggers the death of damaged cells^6^, which occurs in neurodegenerative disorders including Parkinson’s disease (PD) and Alzheimer’s disease (AD)^1^. *In vitr*o and *in vivo* studies indicate that UPR activation is a “double-edged sword”, as evidence suggests that short-term activation plays a protective role whereas long-term activation results in synaptic failure, impaired synaptic plasticity, and ultimately, cell death^7, 8^. Together, these studies reinforce the idea that proper ER proteostasis is key for sustaining neuronal connectivity and function.

The functional link between ER stress, the UPR, and neurodegeneration has been extensively investigated over the last decade^9^. Activation of the UPR has been reported in *post mortem* brain tissues from patients with a number of neurodegenerative disorders. Increases in markers of UPR activation, in particular, the increased phosphorylation of PERK and eIF2α, have been observed in PD, AD, and other tauopathies^9, 10^. Selective neuronal populations seem to be especially vulnerable to ER stress^11^, so it is perhaps unsurprising that the clinical manifestation of neurodegenerative diseases is initiated by the selective alteration in the function of distinct neuronal populations. Activation of PERK-eIF2α signaling and a co-localization of phosphorylated PERK and a-synuclein, a disease-specific misfolded protein, have been reported in dopaminergic (DA) neurons of the substantia nigra in brain tissue of PD patients^10, 12^. Beyond its critical role in the control of voluntary movement via the nigrostriatal pathway, DA signaling contributes to synaptic plasticity underlying learning and memory in specific brain regions, including the hippocampus, amygdala, and prefrontal cortex, and altered DA modulation affects the encoding and maintenance of memories^13–15^.

The findings described above suggest that modulation of the UPR in specific subpopulations of neurons may supply therapeutic benefits in the treatment of neurodegenerative disorders. Consistent with this idea, genetic and pharmacological modulation of PERK-eIF2α signaling has emerged as a therapeutic target in neurodegenerative diseases, primarily due to its role in mediating eIF2-dependent protein synthesis^16–18^. However, these studies have proved to be as controversial as promising ^19, 20^, indicating a new layer of complexity to the involvement of UPR in neurodegeneration. In the current study, we explored the cell type-specific modulation of PERK-eIF2α signaling in DA neurons. We used genetic approaches that included Cre-Lox recombination technology to selectively delete PERK in DA neurons of both the nigrostriatal and mesocorticolimbic pathways to investigate the consequences of manipulating the UPR on motor and cognitive function. Notably, we found that genetic disruption of PERK-eIF2α signaling in DA neurons in mice resulted in multiple age-dependent motor and cognitive phenotypes. In addition, *de novo* translation studies revealed dysregulated protein synthesis in DA neurons, and fast-scan cyclic voltammetry (FSCV) in *ex vivo* striatal slices showed an age-dependent alteration in DA release and DAT-mediated uptake that contribute to the behavioral phenotypes caused by the deletion of PERK in DA neurons. Overall, our findings show that proper cell type-specific regulation of PERK-eIF2 signaling in DA neurons is required for normal motor and cognitive function, and that the UPR plays a critical role in maintaining DA neuron function during development and aging. These findings highlight the need for careful evaluation of the effects of sustained PERK inhibition in neurons, which has been proposed as a therapeutic strategy in the treatment of several neurodegenerative disorders.

## Results

### Conditional deletion of PERK in DA neurons leads to multiple age-dependent motor and cognitive phenotypes in mice

In order to evaluate whether proper cell type-specific regulation of PERK-eIF2α signaling in DA neurons is required for normal cognitive and motor function, we generated mice containing a DA transporter (DAT) promoter-driven *Cre* transgene (DAT-Cre; Jackson Laboratory, stock number: 006660)^21^ and a conditional allele of *Perk* (*Perk*^loxP^; termed PERK^f/f^; **Fig. 1a**)^22^. The expression of the *Cre* transgene and the *Perk*^loxP^ allele was determined using PCR-specific primers (**Fig. 1b**). The resulting conditional knockout mice (PERK^f/f^ DAT-Cre), which lack PERK in DA neurons of both the ventral tegmental area (VTA) and the substantia nigra pars compacta (SNc) represent the primary experimental mouse line used here, along with their littermate control mice (WT DAT-Cre).

**Figure 1.**
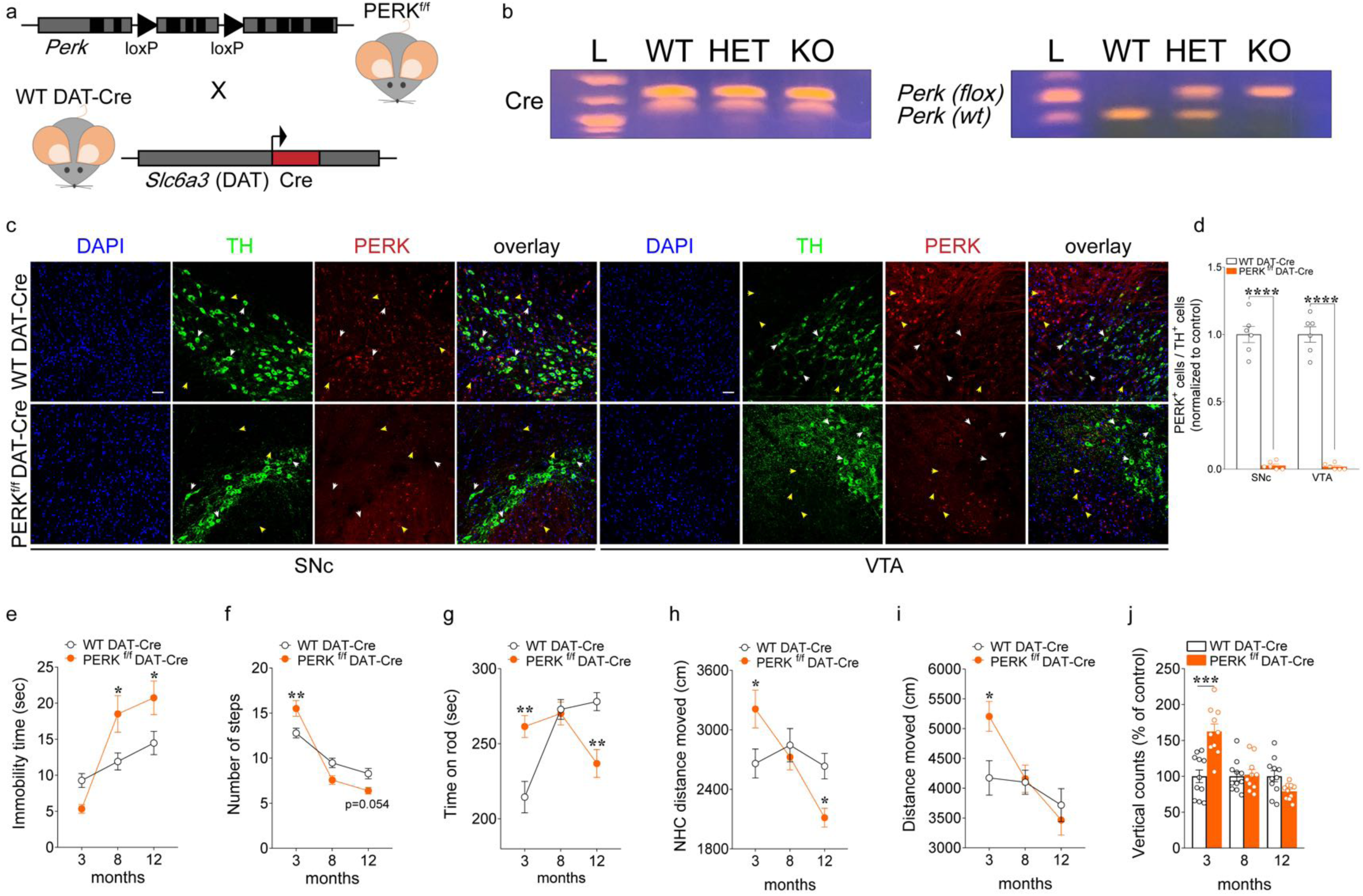
Deletion of PERK from DA neurons induces motor facilitation at early ages, but worsens motor performance in older mice. (**a**) Schematic representation of DAT-neuron specific deletion of PERK in PERK^f/f^ mice crossed with WT DAT-Cre mice. (**b**) PCR identification of alleles of Perk^loxP^ and DAT-driven Cre. (**c**) Immunofluorescent detection of TH^+^ (green) neurons and PERK (red) expression in SNc and VTA DA neurons of PERK^f/f^ DAT-Cre and WT DAT-Cre mice, confirming positive targeting of dopaminergic (TH^+^) neurons for the deletion of PERK (scale bars represent 50 µm). White arrows indicate dopaminergic neurons (green) and PERK (red) co-staining; yellow arrows indicate non dopaminergic neurons and PERK (red) staining (**d**) Summary plot showing the ratio of TH+ cells in both the SNc and the VTA that co-labeled for PERK in PERK^f/f^ DAT-Cre vs. WT DAT-Cre control mice (n = 6 mice each, unpaired t test, SNc: t*_(10)_*= 15.85, *P* <0,0001; VTA: t*_(10)_*= 16.86, *P* <0,0001) after treatment with thapsigargin. PERK^f/f^ DAT-Cre and their WT DAT-Cre littermates mice were subjected to a set of tests, including the bar (**e**), drag (**f**), rotarod (**g**), novel home cage (**h**), and open field (**i,j**) tests to investigate locomotor activity, from 3 up to 12 months of age. (**e**) Summary plot of immobility time (sec) during bar test in PERK^f/f^ DAT-Cre *versus* WT DAT-Cre mice (two-way RM ANOVA, time x genotype, F*_(2, 93)_* = 6.80, *P* < 0.01). (**f**) Summary plot of average number of steps during drag test (two-way RM ANOVA, time x genotype, F*_(2, 93)_* = 10.31, *P* < 0.001). (**g**) Summary plot of latency to fall from the rotating rod measured as average of two days (4 trials/day) test (two-way RM ANOVA, time x genotype, F*_(2, 44)_* = 14.93, *P* < 0.001). (**h**) Summary plot of the novelty-induced locomotor activity expressed as a novel home cage (NHC) distance moved (cm) in the first 10 minutes interval of a 60 minutes test during novel home cage test (two-way RM ANOVA, time x genotype, F*_(2, 58)_* = 6.52, *P* < 0.01). (**i,j**) Summary plot of (**i**) spontaneous locomotor activity expressed as distance moved (cm) and (**j**) vertical activity (number of counts) during the open field test over 15 min (two-way RM ANOVA, time x genotype; **i**, F*_(2, 38)_* = 4.18, *P* < 0.05; **j**, F*_(2, 38)_* = 18.89, *P* < 0.001). Mice were analyzed using two-way RM ANOVA followed by the Bonferroni’s test for multiple comparisons. \**P* < 0.05, \*\**P* < 0.01, \*\*\**P* < 0.001 different from age-matched littermates. All data are shown as mean ± s.e.m. of *n* = 18-15, *n* = 17-13 and *n* = 19-17 mice/genotype at 3, 8 and 12 months of age, respectively (**e,f**); *n* = 12 mice/genotype (**g**); *n* = 15-16 mice/genotype (**h**) *n* = 10-11 mice/genotype (**i,j**).

Cell-specific deletion of PERK in DA neurons was first verified at the protein expression level by treating coronal midbrain slices containing the VTA and SNc with thapsigargin, which inhibits ER Ca^2+^ sequestration and is a potent inducer of ER stress and eIF2α phosphorylation. Immunostaining for PERK after thapsigargin exposure was clearly seen in tyrosine-hydroxylase positive (TH+) DA neurons in both SNc and VTA in WT DAT-Cre mice, whereas PERK staining was not detected in SNc or VTA DA neurons of PERK^f/f^ DAT-Cre mice, although PERK expression in non-DA (TH-) cells remained intact (**Fig. 1c-d**). Consistent with these results, downstream UPR targets of PERK such as p-eIF2α (**Supplementary Fig. 1a-d**) and ATF4 (**Supplementary Fig. 1e-f**) were significantly reduced in both SNc and VTA DA neurons of PERK^f/f^ DAT-Cre mice compared to thapsigargin-treated controls. Taken together, these results demonstrate a reduced UPR after thapsigargin-induced ER-stress in midbrain DA neurons of PERK^f/f^ DAT-Cre mice, confirming positive targeting of DA neurons for the deletion of PERK.

To provide further verification of the quality of the recombination system and the cell-specificity of *Cre* transgene expression in DAT^+^ neurons, we generated a separate mouse line containing the DAT-Cre transgene, the *Perk*^loxP^ allele and Ai14, a *Cre* reporter allele that has a *loxP*-flanked STOP cassette preventing transcription of a CAG promoter-driven red fluorescent protein variant (tdTomato), all inserted into the *Gt(ROSA)26Sor* locus (B6; 129S6- *Gt(ROSA)26Sor^tm^*^14(CAG–tdTomato)^*^Hze^*; Jackson Laboratory, stock number: 007914)^23^ that express tdTomato fluorescence following Cre-mediated recombination (**Supplementary Fig. 2a**). Thus, along with the deletion of PERK from in neurons in which Cre is expressed (DAT^+^ neurons), the STOP cassette is removed and the tdTomato protein is expressed. We found a complete overlap between cells expressing tdTomato fluorescence (Cre-reporter expression) and the DA neuronal marker tyrosine hydroxylase (TH; **Supplementary Fig. 2b**) in both PERK^f/f^/CAG^floxStop-tdTomato^DAT-Cre and wild-type (WT)/CAG^floxStop-tdTomato^DAT-Cre mice. Moreover, we found no staining for PERK in tdTomato co-stained neurons (**Supplementary Fig. 2c,e**) of PERK^f/f^/CAG^floxStop-tdTomato^DAT-Cre mice, confirming the specificity of the Cre-recombinase system for the DA neurons. Along with these results, we detected a significant reduction in p-eIF2α levels in DA neurons in both SNc and VTA of PERK^f/f^/CAG^floxStop-tdTomato^DAT-Cre mice (**Supplementary Fig. 2d,f**).

To determine consequences of the sustained disruption of PERK-eIF2α signaling in DA neurons on motor ability in mice across ages, we examined PERK^f/f^ DAT-Cre mice and their age-matched WT DAT-Cre littermate controls in a series of behavioral tests at 3, 8, and 12 months of age. Mice were tested for different motor skills, including akinesia (bar test), bradykinesia (drag test), general motor activity (rotarod test), locomotor activity induced by novelty (NHC), and spontaneous horizontal and vertical locomotor activity in the open field (OF). PERK^f/f^ DAT-Cre mice displayed a striking biphasic change in motor performance as the mice aged. Specifically, compared to age-matched controls, mice lacking PERK in DA neurons generally exhibited a hyperactive motor phenotype at 3 months of age, had motor behavior that was similar to controls at 8 months, but then showed reversal to a hypoactive phenotype at 12 months of age (**Fig. 1e-j**). In the bar test, young PERK^f/f^ DAT-Cre mice showed a trend for reduced immobility time *versus* WT DAT-Cre, whereas 8 and 12-month old PERK^f/f^ DAT-Cre mice showed increased akinesia compared to WT DAT-Cre mice (**Fig. 1e**). Consistent with a hyperactive phenotype at 3 months, young PERK^f/f^ DAT-Cre mice showed significantly greater stepping activity than WT DAT-Cre mice in the drag test (**Fig. 1f**). Although overall stepping activity decreased with age for both genotypes in this test, the performance of PERK^f/f^ DAT-Cre mice exhibited a more pronounced decline at 12 months compared to control mice. Motor facilitation was also seen in the rotarod test for young PERK mutant mice, with significantly enhanced motor skill acquisition compared to WT DAT-Cre mice at 3 months (**Fig. 1g**). This motor difference was normalized at 8 months, then reversed at 12 months, with a marked drop in rotorod performance in PERK^f/f^ DAT-Cre compared to that of age-matched controls (**Fig. 1g**). Novelty-induced (**Fig. 1h**) and spontaneous (**Fig. 1i**) horizontal locomotor activity followed similar patterns, with hyperactivity motor phenotype in PERK^f/f^ DAT-Cre mice at 3 months, but a significant decline associated with aging in DAT-Cre *versus* WT at 12 months (**Fig. 1h,i**). Moreover, PERK^f/f^ DAT-Cre mice displayed a significantly enhanced vertical locomotor activity in the OF arena compared with their controls at 3 months of age, although the difference was lost as the mice aged (**Fig. 1j**).

PERK is a key regulator of eIF2-dependent translation, which is a key molecular process underlying learning and memory formation ^24–26^. To determine whether the conditional deletion of PERK in DA neurons impacts cognitive, as well as motor function, we examined 3- and 12-month old PERK^f/f^ DAT-Cre mice and age-matched littermate WT DAT-Cre mice in a series of behavioral tasks to test learning and memory. First, we tested spatial learning and memory in 3-month old mice in the Morris water maze (MWM), a hippocampus-dependent water escape task. During the acquisition of the hidden platform phase of the water maze task, both genotypes showed a day-to-day decrease in escape latency, although the daily improvement was less in PERK^f/f^ DAT-Cre mice which consistently exhibited longer escape latencies than WT DAT-Cre controls (**Fig. 2a**). In the probe test when the platform was removed, PERK^f/f^ DAT-Cre mice spent less time in the target quadrant (**Fig. 2b**) and crossed the platform location fewer times (**Fig. 2c**) compared to WT DAT-Cre mice, with representative swim paths shown in **Fig. 2d**. Notably, 12-month old PERK^f/f^ DAT-Cre mice displayed learning and memory deficits that paralleled those seen in the 3-month old mice, including longer escape latencies during training (**Fig. 2h**), reduced time in the target quadrant (**Fig. 2i**), and fewer crossings of the platform location (**Fig. 2j**) during the probe test compared to their WT DAT-Cre counterparts. The impaired performance seen in mice lacking PERK in midbrain DA neurons suggest that the sustained disruption of PERK-eIF2α signaling alters a DA-dependent contribution to this hippocampus-dependent spatial memory task.

**Figure 2.**
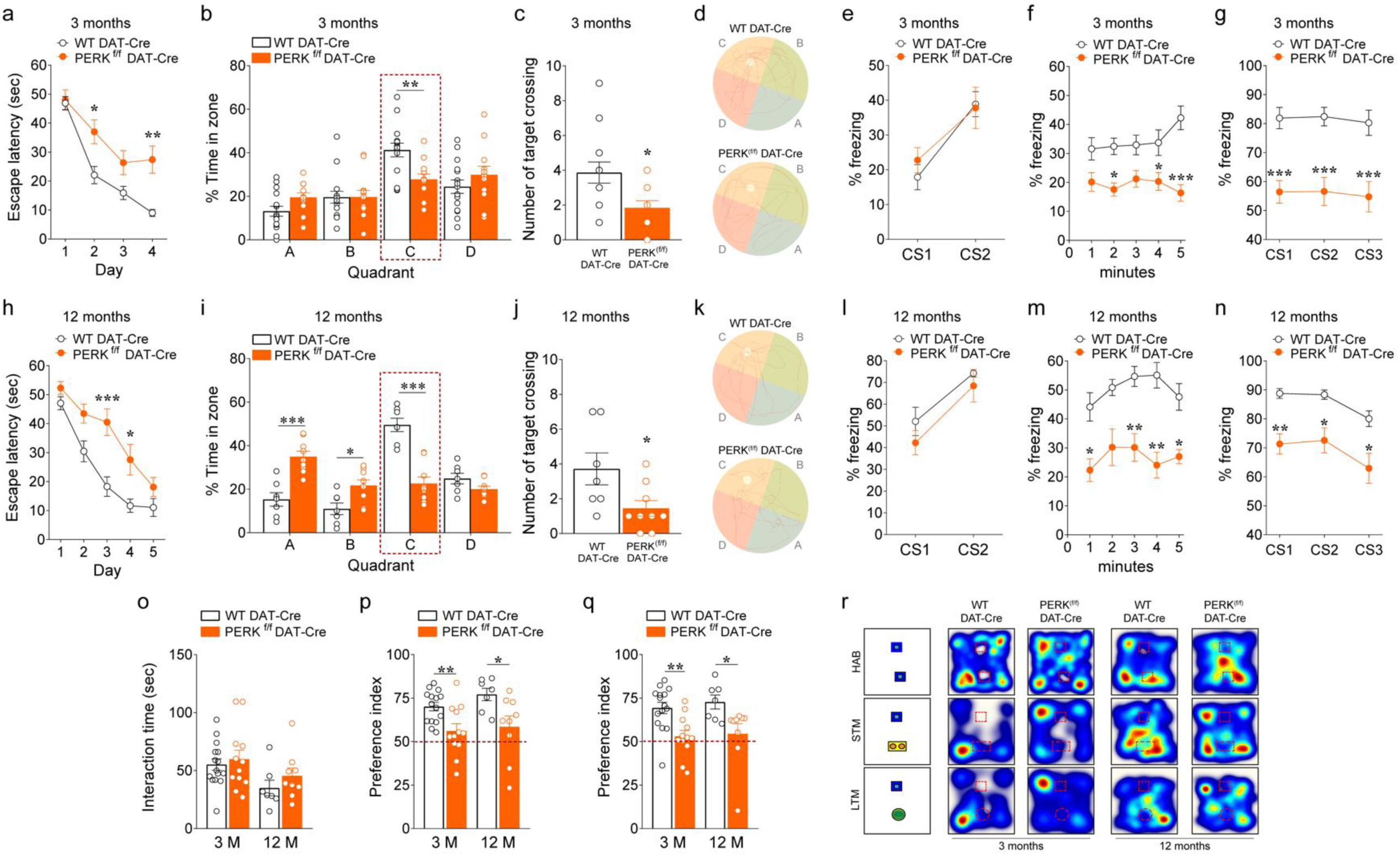
Deletion of PERK from DA neurons results in multiple cognitive phenotypes in mice. (**a-c**) Summary plots of (**a**) average latency to find the hidden platform during a 4-day training protocol, (**b**) percentage time spent in each zone, and (**c**) average number of times crossing the location of the previously hidden platform during probe test in 3- month old PERK^f/f^ DAT-Cre *versus* WT DAT-Cre mice in the MWM test (**a**, two-way RM ANOVA, followed by Bonferroni’s multiple comparisons test, time x genotype, F*_(3, 75)_* = 3.53, *P* < 0.05; **b**, two-way RM ANOVA, followed by Bonferroni’s multiple comparisons test, quadrant x genotype, F*_(3, 75)_* = 3.77, *P* < 0.05; **c**, unpaired *t* test, t_(25)_ = 2.607, *P* = 0.02; *n* = 12 PERK^f/f^ DAT-Cre mice; *n* = 15 WT DAT-Cre mice). (**d**) Representative swim paths in the MWM test. (**e-g**) Summary plots of average percentage of freezing during (**e**) training, (**f**) exposure to the context 24 hours after training, and (**g**) exposure to 3 CS presentations in a novel context in the associative threat memory test in 3-month old PERK^f/f^ DAT-Cre *versus* WT DAT-Cre mice (**e**, two-way RM ANOVA, followed by Bonferroni’s multiple comparisons test, time x genotype, F*_(1,25)_* = 0.76, *P* = 0.39; **f**, two-way RM ANOVA, followed by Bonferroni’s multiple comparisons test, genotype, F*_(1, 25)_* = 20.45, *P* = 0.0001; **g**, two-way RM ANOVA, followed by Bonferroni’s multiple comparisons test, genotype, F*_(1, 25)_* = 28.53 *P* <0.0001; *n* = 12 PERK^f/f^ DAT-Cre mice; *n* = 15 WT DAT-Cre mice). (**h-j**) Summary plots of (**h**) average latency to find the hidden platform during a 4-day training protocol, (**i**) percentage time spent in each zone and (**j**) average number of times crossing the location of the previously hidden platform during probe test in 12- month old PERK^f/f^ DAT-Cre *versus* WT DAT-Cre mice in the MWM test (**h**, two-way RM ANOVA, followed by Bonferroni’s multiple comparisons test, time x genotype, F*_(4, 56)_* = 2.61, *P* < 0.05; **i**, two-way RM ANOVA, followed by Bonferroni’s multiple comparisons test, quadrant x genotype, F*_(3, 42)_* = 22.84, *P* < 0.001; **j**, unpaired *t* test, t_(14)_ = 2.392, *P* = 0.03; *n* = 9 PERK^f/f^ DAT-Cre mice; *n* = 7 WT DAT-Cre mice). (**k**) Representative swim paths in the MWM test. (**l-n**) Summary plots of average percentage of freezing during (**l**) training, (**m**) exposure to the context 24 hours after training, and (**n**) exposure to3 CS presentations in a novel context in the associative threat memory test in 12-month old PERK^f/f^ DAT-Cre *versus* WT DAT-Cre mice (**l**, two-way RM ANOVA, followed by Bonferroni’s multiple comparisons test, time x genotype, F*_(1,14)_* = 0.19, *P* = 0.67; **m**, two-way RM ANOVA, followed by Bonferroni’s multiple comparisons test, genotype, F*_(1, 14)_* = 31.13, *P* < 0.0001; **n**, two-way RM ANOVA, followed by Bonferroni’s multiple comparisons test, genotype, F*_(1, 14)_* = 22.98 *P* <0.001; *n* = 9 PERK^f/f^ DAT-Cre mice; *n* = 7 WT DAT-Cre mice). (**o-q**) Summary plots of (**o**) interaction time with familiar objects, and (**p,q**) preference indices of mice towards a novel object introduced in the novel object recognition test in 3- month and 12-month old PERK^f/f^ DAT-Cre *versus* WT DAT-Cre mice (unpaired *t* test; **o**, t_(25)_ = 0.513, *P* = 0.61 and t_(14)_ = 1.074, *P* = 0.30 respectively in 3-month and 12-months old mice; **p**, t_(25)_ = 3.079, *P* < 0.01 and t_(14)_ = 2.417, *P* < 0.05 respectively in 3-month and 12- months old mice; **q**, t_(25)_ = 3.433, *P* < 0.01 and t_(14)_ = 2.427, *P* < 0.05 respectively in 3-month and 12-months old mice; 3-month old mice: *n* = 12 PERK^f/f^ DAT-Cre mice, *n* = 15 WT DAT-Cre mice; 12-month old mice: *n* = 9 PERK^f/f^ DAT-Cre mice, *n* = 7 WT DAT-Cre mice). (**r**) Representative heat maps of the interaction with the objects for each genotype in the novel object recognition test. All data are shown as mean ± s.e.m. \**P* < 0.05, \*\**P* < 0.01 and \*\*\**P* < 0.001 PERK^f/f^ DAT-Cre *versus* WT DAT-Cre mice.

To confirm our findings that PERK-eIF2α signaling disruption in DA neurons impacts learning and memory, we tested PERK^f/f^ DAT-Cre mice and their WT DAT-Cre littermates on two additional tasks: novel object recognition and an associative threat memory task (**Fig. 2**). In the novel object recognition task, we found that 3-month old PERK mutant mice exhibited similar interactions with familiar objects during training to age-matched WT DAT-Cre mice (**Fig. 2o**), but showed a decreased preference for the novel object, indicating a significantly impaired short-term memory (STM) performance during the test *versus* controls (**Fig. 2p**). Moreover, young PERK^f/f^ DAT-Cre mice also demonstrated a significantly reduced preference for the novel object *versus* controls when long-term memory (LTM) was examined 24 hours after the test (**Fig. 2q**). Representative heat maps are shown in **Fig. 2r**. Again, 12-month old PERK^f/f^ DAT-Cre mice recapitulated the phenotype exhibited by the 3-month old mice, with impaired STM (**Fig. 2p**) and LTM (**Fig. 2q**) during the novel object recognition task (**Fig. 2o-r**). Combined, these results suggest that PERK in DA neurons is important for frontal and temporal cortex-dependent sensory information processing. Similar to the novel object recognition task, 3-month old PERK^f/f^ DAT-Cre mice exhibited altered associative learning and memory when tested with a threat memory task (**Fig. 2e-g**). Both genotypes performed similarly during training, showing a similar freezing behavior (**Fig. 2e**). However, PERK^f/f^ DAT-Cre mice displayed significantly reduced freezing 24 hours after training for both the context (**Fig. 2f**) and the cue (**Fig. 2g**) compared to their WT DAT-Cre littermates. Similarly, 12-month old PERK^f/f^ DAT-Cre mice showed no difference in freezing compared to the WT DAT-Cre mice during training (**Fig. 2l**), but displayed a significant decrease in freezing time to both the context (**Fig. 2m**) and the cue (**Fig. 2n**). These behavioral studies indicate that the deletion of PERK in DA neurons not only affects hippocampal and frontal cortex-dependent function, but also results in amygdala-dependent memory impairments, suggesting functional alterations of the mesocorticolimbic pathways^27^.

### Deletion of PERK in DA neurons alters *de novo* translation and multiple behaviors via eIF2α signaling disruption

As mentioned above, phosphorylation of eIF2α on serine 51 is a mechanism through which PERK downregulates global protein synthesis under a variety of cellular stress conditions. Consistent with blocked expression of PERK, the phosphorylation of eIF2α was reduced in DA neurons of 3-month old PERK^f/f^ DAT-Cre mice (**Supplementary Fig. 2d**). Given that eIF2α phosphorylation is a key step in translational control under normal physiological conditions and during ER-stress, we investigated the effect of removing PERK on *de novo* protein synthesis in DA neurons (**Fig 3**). Coronal midbrain slices containing SNc and VTA DA neurons were subjected to fluorescent non-canonical amino acid tagging (FUNCAT) of newly synthesized proteins (**Fig. 3a**). We observed an increase in *de novo* translation in DA neurons of the SNc of PERK^f/f^ DAT-Cre mice compared to wild-type DAT-Cre littermates (**Fig. 3 b,c**). Similar to SNc, VTA DA neurons of PERK^f/f^ DAT-Cre mice also exhibited a net increase in newly synthesized proteins (**Fig. 3b,d**).

**Figure 3.**
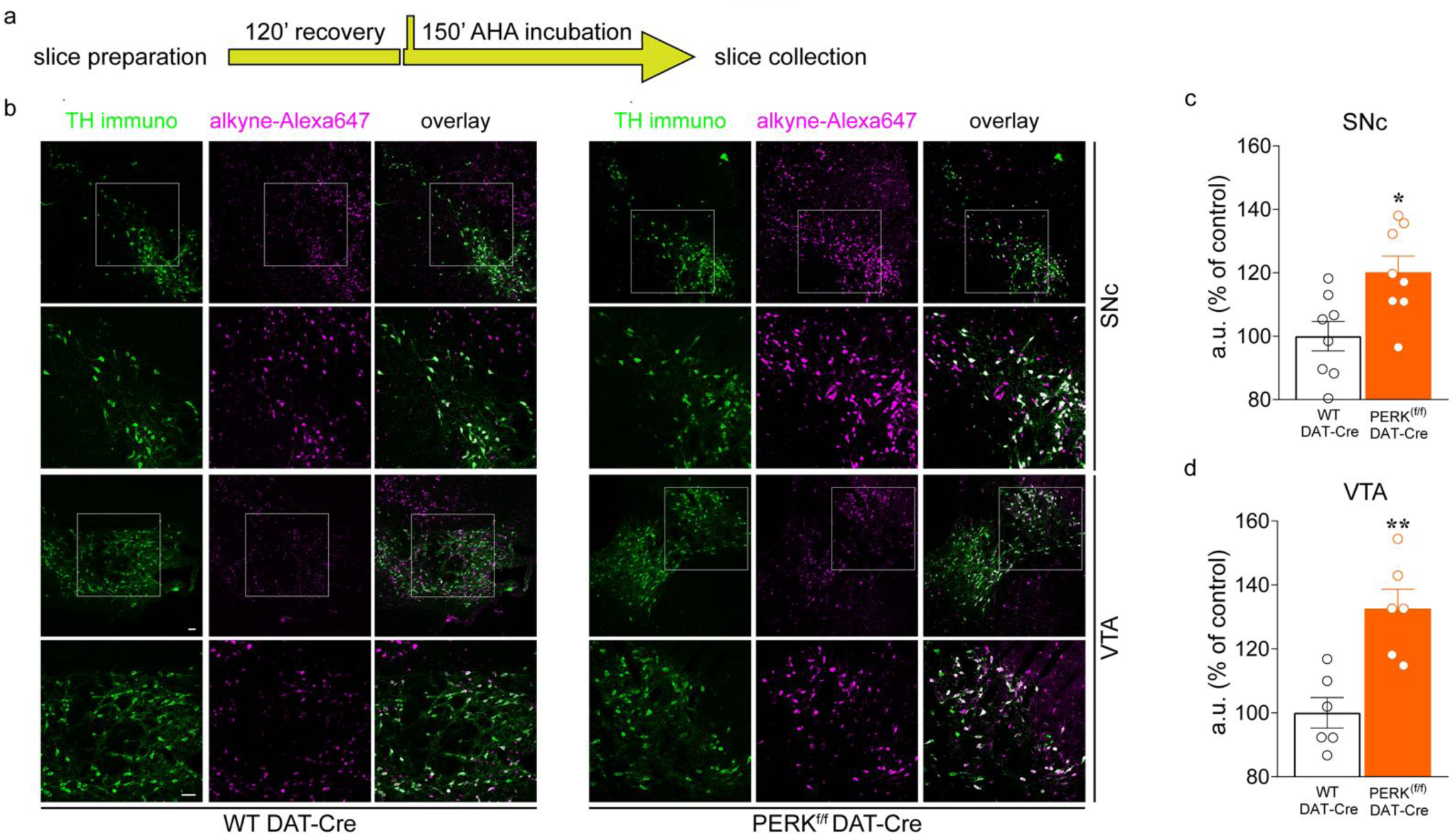
Deletion of PERK in DA neurons causes dysregulated *de novo* translation in mice. (**a**) Schematic for FUNCAT experiments. (**b**) Representative immunofluorescence images of TH-IR (green) and incorporation of AHA (magenta) detected by FUNCAT with alkyne-Alexa 647 in SNc and VTA DA neurons from midbrain coronal slices of 3-month old PERK^f/f^ DAT-Cre mice and their WT DAT-Cre littermates. Insets in the first and third rows are magnified in the second and fourth rows, respectively (scale bars represent 50 µm). (**c,d**) Quantification of increased AHA-alkyne-Alexa 647 signal in fluorescent arbitrary units (a.u.) expressed as % of control in TH^+^ neurons (Green) from SNc (**c**) and VTA (**d**) of PERK^f/f^ DAT-Cre *vs*. WT DAT-Cre mice. Cell soma intensity was measured in ImageJ. Statistical significance was determined by using Student’s *t* test (PERK^f/f^ DAT-Cre *vs*. WT DAT-Cre mice; unpaired *t* test; **c,** t_(14)_ = 2.933, *P* = 0.011; **d**, t_(10)_ = 4.215, *P* = 0.002). Data are shown as mean ± s.e.m. of *n* = 6/8 mice per group (average of *n* = 40 somas per slice, *n* = 2 slices per mouse, from three independent experiments) \**P* < 0.05, \*\**P* < 0.01.

To confirm that the behavioral results obtained by deleting PERK in DA neurons are due to decreased eIF2α phosphorylation and not due to other cellular functions of PERK, we bred DAT-Cre mutant mice (Jackson Laboratory, stock number: 006660)^21^ with conditional phospho-mutant eIF2α mice^28^ (knock-in eIF2α^(S51A/ S51A)^ mice), where the serine 51 residue has been mutated to alanine (S51A; **Fig. 4a**). As a result, both the *floxed* wild-type and the mutated eIF2α gene were expressed in all cells of eIF2α^(S51A/S51A)^ DAT-Cre mice, except for the DA neurons (DAT^+^ neurons), where only the mutated eIF2α^(S51A/S51A)^ form is expressed (see Methods section for details). The expression of the *Cre* transgene and the eIF2α^(S51A/S51A)^ allele was determined using PCR-specific primers (**Fig. 4b**). Then, we verified the cell-specificity of the *Cre* system by treating coronal slices containing the midbrain with thapsigargin to induce ER stress and eIF2α phosphorylation. Levels of p-eIF2α detected by immunofluorescence were significantly reduced (∼90%) in both SNc and VTA DA neurons of eIF2α^(S51A/S51A)^ DAT-Cre mice compared to thapsigargin-treated controls (**Fig. 4c-d**). These data also are consistent with western blotting results showing significantly lower levels of p-eIF2α in both VTA and SNc of eIF2α^(S51A/S51A)^ DAT-Cre mice (**Fig. 4e,j**). As mentioned above, phosphorylation eIF2α on serine 51 is a mechanism through which PERK downregulates global protein synthesis under a variety of cellular stress conditions. To further confirm the cell-specificity expression of the phospho-mutant eIF2α (eIF2α^(S51A/S51A)^) in DA neurons, we investigated *de nov*o translation in DA neuron from thapsigargin-treated midbrain coronal slices with surface sensing of translation (SUnSET). We found a robust increase in *de novo* protein synthesis in both VTA (∼50%; **Fig. 4f**) and SNc (∼80%; **Fig. 4k**) DA neurons of eIF2α^(S51A/S51A)^ DAT-Cre mice compared to controls. Collectively, these data confirm positive targeting of DA (DAT^+^) neurons for the expression of eIF2α^(S51A/S51A)^ and the disruption of the eIF2α translational control pathway.

**Figure 4.**
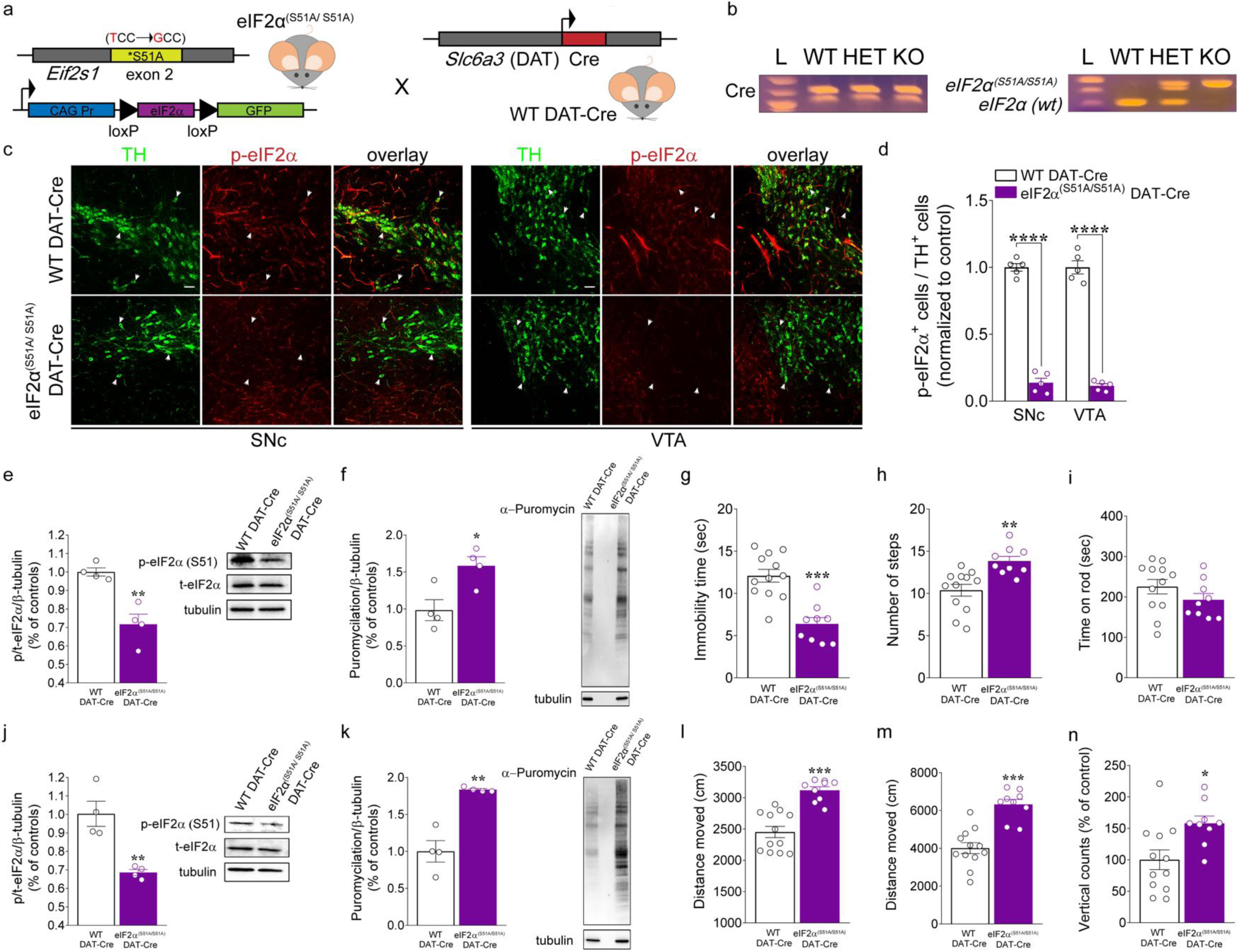
eIF2α^(S51A/S51A)^ DAT-Cre mice display enhanced locomotor activity. (**a**) Schematic representation of DAT-neuron specific expression of phospho-mutant eIF2α in eIF2α^(S51A/S51A)^ mice crossed with WT DAT-Cre mice. (**b**) PCR identification of alleles of eIF2α^(S51A/S51A)^ and DAT-driven Cre. (**c**) Immunofluorescent detection of TH^+^ (green) neurons and phospho-eIF2α (p-eIF2α; red) expression in SNc and VTA DA neurons of eIF2α^(S51A/S51A)^ DAT-Cre and WT DAT-Cre mice, confirming positive targeting of dopaminergic (TH^+^) neurons for the expression of phospho-mutant eIF2α (scale bars represent 50 µm). Arrows indicate dopaminergic neurons (green) and p-eIF2α (red) co-stain. (**d**) Summary data showing the ratio of TH+ cells in both the SNc and the VTA that co-labeled for p-eIF2α in eIF2α^(S51A/S51A)^ DAT-Cre vs. WT DAT-Cre control mice (*n* = 5 mice each, unpaired t test, SNc: t*_(8)_*= 19.61, *P* <0,0001; VTA: t*_(8)_*= 17.19, *P* <0,0001). (**e,j**) Representative western blots (right panel) and quantification of phosphorylation of eIF2α in VTA (**e**) and SNc (**j**) of eIF2α^(S51A/S51A)^ DAT-Cre and WT DAT-Cre mice. Summary plot showed a robust decrease in the phosphorylation of eIF2α in both VTA (**e**; unpaired *t* test, t*_(6)_* = 4.79, *P* = 0.003; *n*= 4 independent lysates from 4 mice per group) and SNc (**j**; unpaired *t* test, t*_(6)_* = 4.54, *P* = 0.004; *n*= 4 independent lysates from 4 mice per group) of eIF2α^(S51A/S51A)^ DAT-Cre mice compared with controls. (**f,k**) Representative western blots (right panel) and quantification of newly synthesized brain proteins in VTA (**f**) and SNc (**k**) of eIF2α^(S51A/S51A)^ DAT-Cre and WT DAT-Cre mice, labelled with puromycin using the SUnSET method (see Methods). Summary plot of puromycilation indicated increased d*e novo* translation in both VTA (**f**; unpaired *t* test, t*_(6)_* = 3.18, *P* = 0.019; *n*= 4 independent lysates from 4 mice per group) and SNc (**k**; unpaired *t* test, t*_(6)_* = 5.72, *P* = 0.0012; *n*= 4 independent lysates from 4 mice per group) of eIF2α^(S51A/S51A)^ DAT-Cre mice compared with controls. 3-month old eIF2α^(S51A/S51A)^ DAT-Cre mice and their WT DAT-Cre littermates mice were subjected to a set of tests including the bar (**g**), drag (**h**), rotarod (**i**), novel home cage, (**l**) and open field (**m,n**) tests to investigate locomotor activity. (**g**) Summary plot of immobility time (sec) during bar test in eIF2α^(S51A/S51A)^ DAT-Cre *versus* WT DAT-Cre mice (unpaired *t* test, t_(19)_ = 5.21, *P* < 0.001). (**h**) Summary plot of average number of steps during drag test (unpaired *t* test, t_(19)_ = 3.67, *P* < 0.01). (**i**) Summary plot of latency to fall from the rotating rod measured as average of two days (4 trials/day) test (unpaired *t* test, t_(19)_ = 1.30, *P* = 0.20). (**l**) Summary plot of the novelty-induced locomotor activity expressed as a distance moved (cm) in the first 10 minutes interval of a 60 minutes test during novel home cage test (unpaired *t* test, t_(19)_ = 5.76, *P* < 0.001). (**m,n**) Summary plot of (**m**) spontaneous locomotor activity expressed as distance moved (cm) and (**n**) vertical activity (number of counts) during the open field test (unpaired *t* test; **m**, t_(19)_ = 2.78 *P* < 0.001; **n**, t_(19)_ = 5.71 *P* = 0.012). All data are shown as mean ± s.e.m. of *n* = 9 eIF2α^(S51A/S51A)^ DAT-Cre mice and *n* = 12 WT DAT-Cre mice. \**P* < 0.05, \*\**P* < 0.01 and \*\*\**P* < 0.001 eIF2α^(S51A/S51A)^ DAT-Cre *versus* WT DAT-Cre mice.

We then used the eIF2α^(S51A/S51A)^ DAT-Cre mice and their WT DAT-Cre littermates to investigate whether the motor phenotypes exhibited by PERK^f/f^ DAT-Cre mice is due to the selective reduction of eIF2α phosphorylation in DA neurons. We tested 3-month old eIF2α^(S51A/S51A)^ DAT-Cre mice and their WT DAT-Cre littermates for different motor skills (**Fig. 4g-I, l-n**). Consistent with the findings with the PERK^f/f^ DAT-Cre mice, we found that conditional phospho-mutant eIF2α mice displayed decreased immobility time in the bar test (**Fig. 4g**) and greater stepping activity in the drag test (**Fig. 4h**) than their WT DAT-Cre counterparts. Although we found no effect of the eIF2α conditional phospho-mutation in the rotarod test (**Fig. 4i**), eIF2α^(S51A/S51A)^ DAT-Cre mice exhibited significantly enhanced horizontal locomotor activity in both the novel home cage (**Fig. 4l**) and open field arena (**Fig. 4m**) tasks, as well as greater vertical locomotor activity compared with WT DAT-Cre littermates (**Fig. 4n**). These findings support the hypothesis that altered general motor ability displayed by the PERK^f/f^ DAT-Cre mice (**Fig. 1**) is due to the disruption of PERK-eIF2α signaling in DA neurons.

To test the hypothesis that PERK regulates learning and memory via eIF2α phosphorylation control in DA neurons, eIF2α^(S51A/S51A)^ DAT-Cre mice were trained in the same cognitive tasks as the PERK^f/f^ DAT-Cre mice. During MWM training, both genotypes showed a day-to-day decrease in escape latency, but the eIF2α^(S51A/S51A)^ DAT-Cre mice spent significantly longer times to locate the hidden platform (**Fig. 5a**). eIF2α^(S51A/S51A)^ DAT-Cre mice not only spent less time in the target quadrant (C; **Fig. 5b**), but also, they exhibited higher preference for the adjacent quadrant (D; **Fig. 5b**) and crossed the platform location fewer times (**Fig. 5c**) than their littermate controls in the probe test. We then tested eIF2α^(S51A/S51A)^ DAT-Cre mice and their littermate controls in the novel object recognition (**Fig. 5h-k**) and associative threat memory tasks (**Fig. 5e-g**). During novel object recognition training, 3-month old conditional phospho-mutant eIF2α mice and their controls exhibited similar interaction profile with familiar objects (**Fig. 5h**). No difference in the preference for the novel object was observed between the genotypes during the short-term memory task (**Fig. 5i**), suggesting intact short-term memory in those mice. However, eIF2α^(S51A/S51A)^ DAT-Cre mice exhibited a reduced preference for the novel object when long-term memory (LTM) was examined 24 hours testing (**Fig. 5j**). Representative heat maps are shown in **Fig. 5k**. The 3-month old conditional phospho-mutant eIF2α mice also had impaired LTM in the associative threat memory task (**Fig. 5e-g**). The eIF2α^(S51A/S51A)^ DAT-Cre mice displayed significantly reduced freezing time during contextual (**Fig. 5f**) and cued (**Fig. 5g**) testing compared with WT DAT-Cre mice. Taken together, these findings are consistent with the cognitive phenotypes exhibited by the PERK^f/f^ DAT-Cre mice and are consistent with the hypothesis that PERK-eIF2α signaling disruption in DA neurons impacts motor functions and multiple cognitive domains in mice.

**Figure 5.**
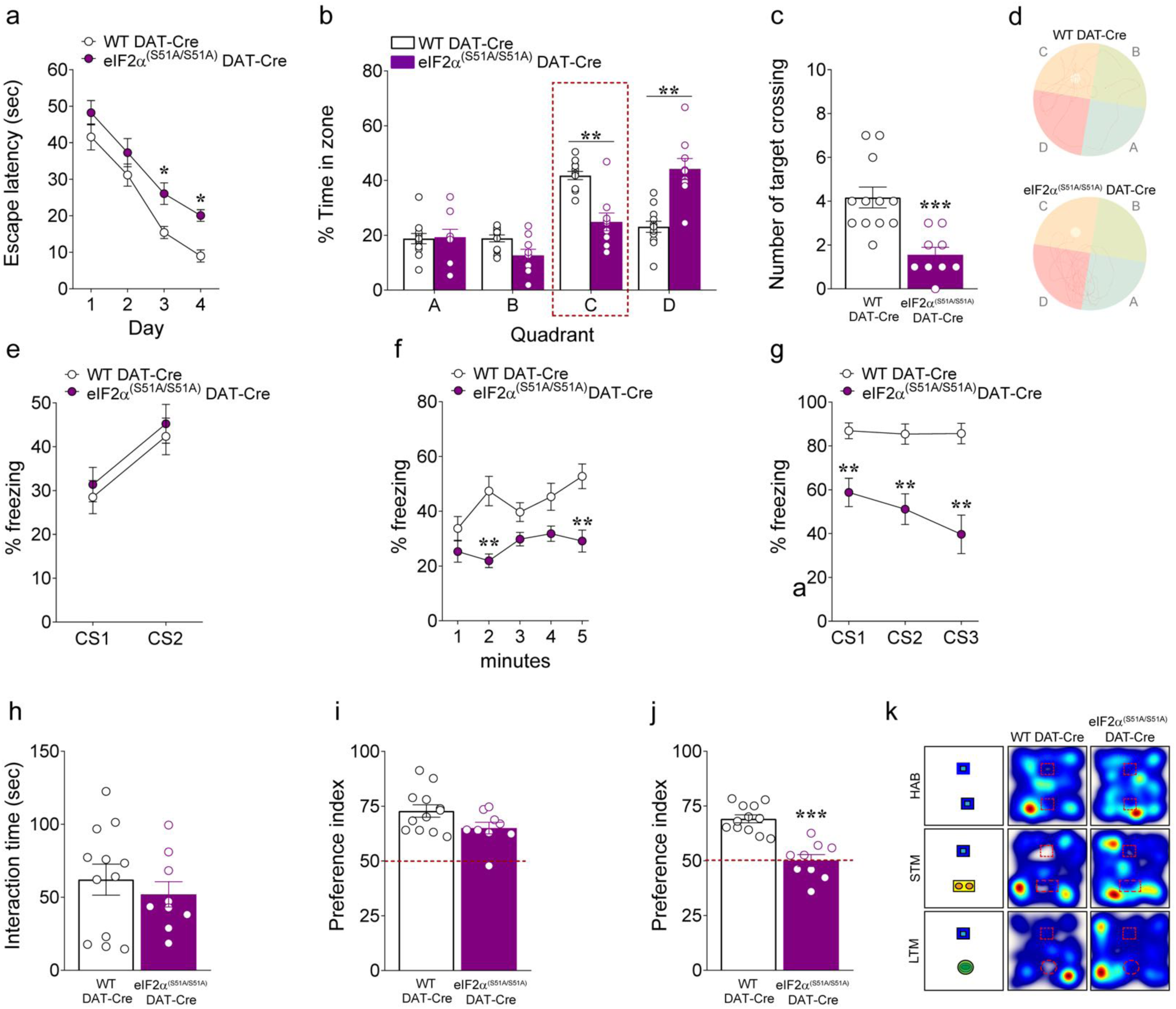
eIF2α^(S51A/S51A)^ DAT-Cre mice display multiple cognitive phenotypes. (**a-c**) Summary plots of (**a**) average latency to find the hidden platform during a 4-day training protocol, (**b**) percentage spent in each zone and (**c**) average number of times spent crossing the location of the previously hidden platform during probe tests in 3-month old eIF2α^(S51A/S51A)^ DAT-Cre *versus* WT DAT-Cre mice in the MWM test (**a**, two-way RM ANOVA, followed by Bonferroni’s multiple comparisons test, time F*_(3, 57)_* = 67.07, *P* < 0.001, genotype F*_(1, 19)_* = 9.49, *P* < 0.01 ; **b**, two-way RM ANOVA, followed by Bonferroni’s multiple comparisons test, quadrant x genotype, F*_(3, 57)_* = 17.65, *P* < 0.001; **c**, unpaired *t* test, t_(19)_ = 4.193, *P* < 0.001). (**d**) Representative swim paths in the MWM test. (**e-g**) Summary plots of average percentage of freezing during (**e**) training, (**f**) exposure to the context 24 hours after training, and (**g**) exposure to 3 CS presentations in a novel context in the associative threat memory test in 3-month old eIF2α^(S51A/S51A)^ DAT-Cre *versus* WT DAT-Cre mice (**e**, two-way RM ANOVA, followed by Bonferroni’s multiple comparisons test, time, F*_(1,19)_* = 78.34, *P* < 0.001; **f**, two-way RM ANOVA, followed by Bonferroni’s multiple comparisons test, genotype, F*_(1, 19)_* = 18.69, *P* < 0.001; **g**, two-way RM ANOVA, followed by Bonferroni’s multiple comparisons test, genotype, F*_(1, 19)_* = 26.76 *P* <0.001). (**h-j**) Summary plots of (**h**) interaction time with familiar objects, and (**i,j**) preference indices of mice towards a novel object introduced in the novel object recognition test in 3-month old eIF2α^(S51A/S51A)^ DAT-Cre *versus* WT DAT-Cre mice (unpaired t test; **h**, t_(19)_= 0.702, *P* = 0.49; **i**, t_(19)_= 1.967, *P* = 0.06; **j**, t_(19)_ = 6.017, *P* < 0.001). (**k**) Representative heat maps of the interaction with the objects for each genotype in the novel object recognition test. All data are shown as mean ± s.e.m. of *n* = 9 eIF2α^(S51A/S51A)^ DAT-Cre mice and *n* = 12 WT DAT-Cre mice. \**P* < 0.05, \*\**P* < 0.01 and \*\*\**P* < 0.001 eIF2α^(S51A/S51A)^ DAT-Cre *versus* WT DAT-Cre mice.

### Deleting PERK in DA neurons leads to a dysregulation of striatal DA release, DAT activity, synaptic plasticity and DA signaling, without affecting DA content

Midbrain DA neurons project to and modulate multiple highly interconnected modules of the basal ganglia, limbic system, and frontal cortex. Alterations in the DA transmission have been reported not only in PD^29^, but also in AD^27^ and Huntington’s disease (HD)^30^, and have been linked to motor as well as cognitive symptoms. Our observation of opposite age-dependent patterns of motor and cognitive changes in PERK^f/f^ DAT-Cre mice led us to hypothesize that the role of PERK in DA neuron function may change as a mouse ages, rather than simply contributing to a degenerative process later in life. As noted, SNc DA neurons project primarily to dStr and play a critical role in motor function and dysfunction via basal ganglia circuitry^29^, whereas VTA DA project to ventral striatum (nucleus accumbens) as well as cortex^14, 31^ and hippocampus^27^. To provide an index of age-dependent effects of selective disruption of the PERK-eIF2α signaling in DA neurons, we used FSCV to quantify single-pulse evoked increases in extracellular DA concentration ([DA]_o_) and DA uptake in *ex vivo* striatal slices from 3-month old and 12-month old PERK^f/f^ DAT-Cre mice and their littermate controls. Data from the NAc serve as a proxy for expected changes in DA release in other VTA-innervated regions, including the hippocampus. Consistent with motor hyperactivity in 3-month old PERK^f/f^ DAT-Cre mice (**Fig. 1e-j**), peak evoked [DA]_o_ in dStr, NAc core, and NAc shell was significantly higher than in littermate controls (**Fig. 6a-d**). In striking contrast, in 12-month old PERK^f/f^ DAT-Cre mice showed significantly lower peak evoked [DA]_o_ compared to control in each region (**Fig. 6e-h**) that was consistent with the motor hypoactivity phenotype seen in PERK mutants at 12 months (**Fig. 1e-j**). Given that net [DA]_o_ reflects both DA release and uptake, we determined maximum DA uptake rate, *V*_max_, from evoked [DA]_o_ records. Increased peak evoked [DA]_o_ could reflect a decrease in *V*_max_, for example. However, we found the opposite, with a significantly higher DA uptake rate in 3-month old PERK^f/f^ DAT-Cre mice in dStr and NAc core (**Fig. 6i-l**; dStr: 3.23 ± 0.12 μM/s PERK^f/f^ DAT-Cre *versus* 2.29 ± 0.13 μM/s in WT DAT-Cre; NAc: 2.28 ± 0.11 μM/s PERK^f/f^ DAT-Cre vs 1.66 ± 0.10 μM/s control). In contrast, *V*_max_ for DA uptake was significantly lower in slices from 12-month-old PERK^f/f^ DAT-Cre mice compared to those from littermate WT DAT-Cre (**Fig. 6i-l**; dStr: 1.84 ± 0.12 μM/s PERK^f/f^ DAT-Cre vs 3.20 ± 0.20 μM/s WT DAT-Cre; NAc: 1.02 ± 0.07 μM/s PERK^f/f^ DAT-Cre vs 2.14 ± 0.10 μM/s WT DAT-Cre).

**Figure 6.**
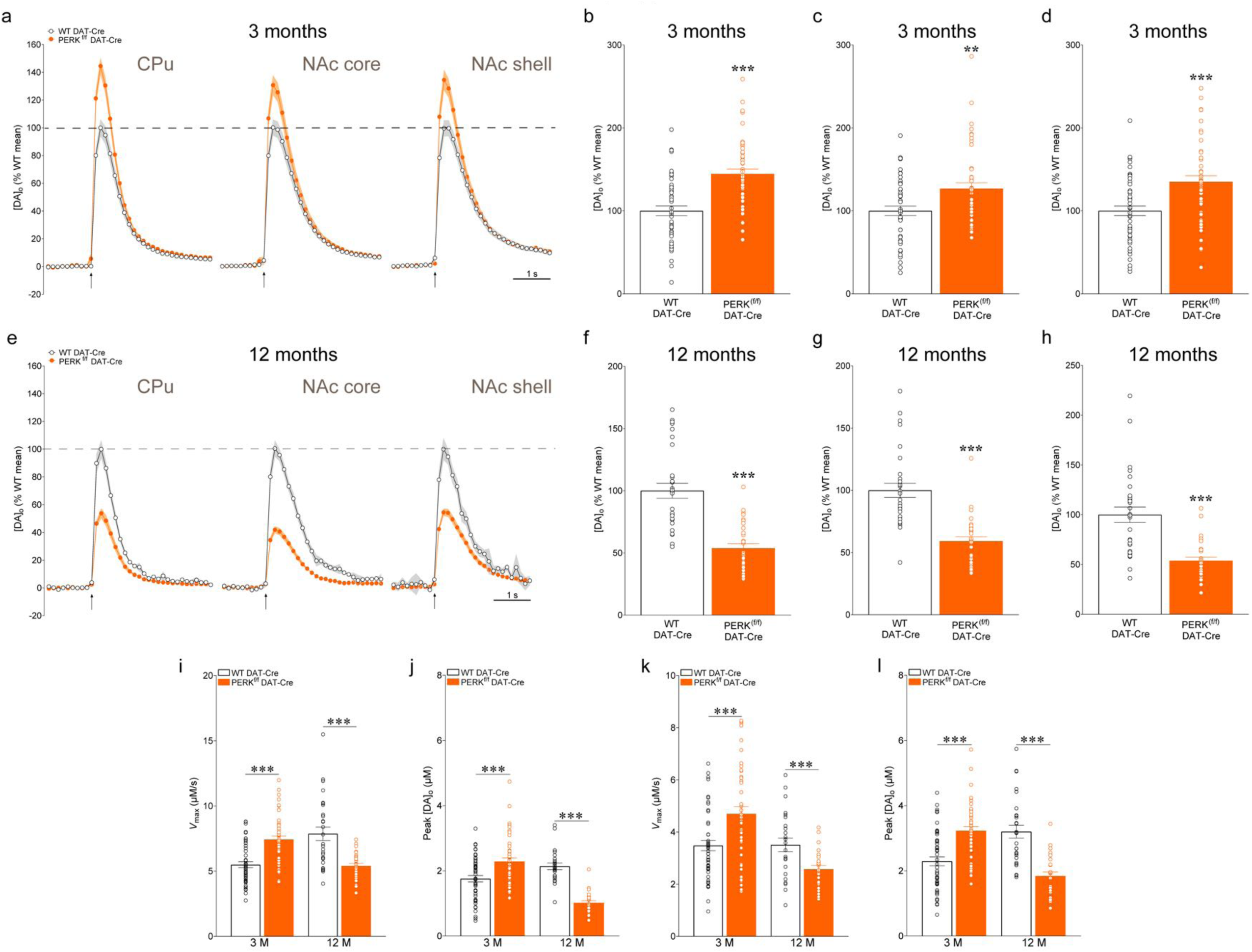
Deletion of PERK in DA neurons alters striatal DA release and DAT activity differently at 3 and 12 months of age in mice. **(a,e)** Average single*-*pulse (1p) evoked [DA]_o_ transients recorded in CPu, NAc core, and NAc shell in PERK^f/f^ DAT-Cre *versus* control DAT-Cre mice at 3 (**a**) and 12 months of age (**e**). (**b-d**) Summary plot of evoked [DA]_o_ peak expressed as % mean control in CPu (unpaired *t*-test, t_(98)_ = 5.345, *P* < 0.001), NAc core (unpaired *t*-test, t_(91)_ = 2.954, *P* < 0.01), and NAc shell (unpaired *t*-test, t_(97)_ = 3.807, *P* < 0.001) in 3-month old mice. (**f-h**) Summary plot of evoked [DA]_o_ peak expressed as % of mean control in CPu (unpaired *t*-test, t_(58)_ = 6.525, *P* < 0.001), NAc core (unpaired *t*-test, t_(58)_ = 6.090, *P* < 0.001), and NAc shell (unpaired *t*-test, t_(58)_ = 5.444, *P* < 0.001) in 12-month old mice. *V*_max_ values (maximum uptake velocity; **i,k**) and *C*_peak_ values (peak concentration; **j,l**) for DAT-mediated DA uptake derived from Michaelis-Menten analysis of single pulse evoked [DA]_o_ records determined using a fixed *K*_m_ of 0.9 µM for each brain region and genotype. (**i**) Summary plot of *V*_max_ values in CPu of WT DAT-Cre *versus* PERK^f/f^ DAT-Cre mice at 3 (unpaired *t*-test, t_(94)_ = 5.589, *P* < 0.001) and 12 (unpaired *t*-test, t_(54)_ = 4.476, *P* < 0.001) months of age. (**j**) Summary plot of *C*_peak_ values in CPu of WT DAT-Cre *versus* PERK^f/f^ DAT-Cre mice at 3 (unpaired *t*-test, t_(94)_ = 5.113, *P* < 0.001) and 12 (unpaired *t*-test, t_(54)_ = 4.476, *P* < 0.001) months of age. (**k**) Summary plot of *V*_max_ values in NAc core of WT DAT *versus* PERK^f/f^ DAT-Cre mice at 3 (unpaired *t*-test, t_(90)_ = 3.616, *P* < 0.001) and 12 (unpaired *t*-test, t_(54)_ = 4.476, *P* < 0.001) months of age. (**l**) Summary plot of *C*_peak_ values in NAc core of WT DAT *versus* PERK^f/f^ DAT-Cre mice at 3 (unpaired *t*-test, t_(90)_ = 3.554, *P* < 0.001) and 12 (unpaired *t*-test, t_(54)_ = 4.476, *P* < 0.001) months of age. Data are means ± s.e.m. of *n* mice, where *n* denotes the number of recording sites sampled from 3 to 5 mice per genotype; \*\**P* < 0.01; \*\*\**P* < 0.001 for WT DAT-Cre *vs.* PERK^f/f^ DAT-Cre mice. Values with R^2^ < 0.95, indicating goodness-of-fit were excluded from the data reported here.

Given that altered [DA]_o_ could reflect changes in DA tissue content, we also determined striatal tissue DA content in the PERK^f/f^ DAT-Cre mice to verify possible changes in DA synthesis that might underlie altered DA availability for release. However, striatal DA levels, quantified using HPLC, did not differ significantly between PERK^f/f^ DAT-Cre mice *versus* their respective controls at either 3 or 12 months (**Supplementary Fig. 3a-c**). Collectively these data show that PERK-eIF2α signaling plays a fundamental role in the dynamic regulation of DA release and uptake as the mice age, independent of changes in DA synthesis.

Although regulation of axonal DA release in the striatum is often linked to the firing patterns of midbrain DA neurons, the activity of striatal cholinergic interneurons (ChIs) can also trigger axonal DA release independently of DA neuron activity via the activation of nicotinic acetylcholine receptors (nAChRs) on DA axons^32, 33^. nAChRs on DA axons are formed by different α and β subunits; previous studies have demonstrated that nAChRs containing β2 subunits mediate ChI-driven DA release^34, 35, 50^. To verify whether the effects of PERK deletion might involve ChIs and altered nAChR-dependent regulation of DA release, we applied dihydro-β-erythroidine (DHβE; 1 μM) a selective antagonist for β2* subunit-containing nAChRs^50^, and again evoked DA release in dStr, NAc core and shell in the same slices recorded under control conditions. The differences in mean peak evoked [DA]_o_ in 3-month and 12-month-old PERK^f/f^ DAT-Cre mice *versus* WT DAT-Cre mice persisted in the presence of DHβE, showing that the effect of PERK deletion in DA neurons on axonal DA release is direct and cell-autonomous, and not an indirect effect of altered regulation by ChIs and nAChR activation (**Supplementary Fig. 4**).

The parallel between age-dependent changes in locomotor activity in PERK^f/f^ DAT-Cre mice (**Fig 1**) and age-dependent changes in evoked DA release (**Fig. 6**) and uptake in these mice strongly suggests that the changes in DA release drive the changes in motor behavior. We therefore tested the DA dependence of the behaviors seen in 3-month old PERK^f/f^ DAT-Cre and WT DAT-Cre. Each genotype was challenged with two different doses (0.01 mg/kg, 0.2 mg/kg; i.p.) of the D1 receptor antagonist SCH 23390. We found that acute SCH 23390 (0.01 mg/kg) injection, which previously was shown to cause motor impairments in naive mice^36^, was sufficient to induce a reduction in locomotor activity in control mice, but was ineffective in PERK^f/f^ DAT-Cre mice, consistent with enhanced DA release in PERK mutants (**Fig. 7a-c**). Similarly, PERK^f/f^ DAT-Cre mice treated with low-dose SCH 23390 showed no differences in their stepping activity (**Fig. 7a**), rotarod performance (**Fig. 7b**), or distance moved (**Fig. 7c**), whereas control mice treated with SCH 23390 (0.01 mg/kg) had reduced locomotor activity (**Fig. 7a-c**). Confirming the DA dependence of motor hyperactivity in PERK^f/f^ DAT-Cre, high-dose SCH 23390 (0.2 mg/kg) reduced locomotor activity in PERK^f/f^ DAT-Cre mice to levels seen in untreated controls (**Fig. 7a-c**). In addition to showing increased DA release at 3 months of age, PERK^f/f^ DAT-Cre mice also exhibit a significant increase in *V*_max_ for DA uptake via the DAT (**Fig. 6i-l**), possibly to compensate for elevated DA release., We challenged 3-month old PERK^f/f^ DAT-Cre mice and their age-matched WT DAT-Cre littermate controls with the DAT inhibitor GBR-12783 (6 mg/kg; 10 mg/kg; i.p.) and examined motor performance. We found that acute DAT blockade with a low dose of GBR-12783 (6 mg/kg) did not change the behavioral responses of the PERK^f/f^ DAT-Cre mice, but significantly increased the locomotor activity of WT DAT-Cre mice, as expected (**Fig. 7d-f**). Inhibition of the DAT by GBR-12783 (6 mg/kg) also increased stepping activity in the drag test (**Fig. 7d**), facilitated rotarod motor performance (**Fig. 7e**) and increased the distance travelled (**Fig. 7f**) in WT DAT-Cre, but not PERK^f/f^ DAT-Cre mice. Consistent with the observed increase in *V*_max_ in PERK mutants, we found that a higher dose of GBR-12783 (10 mg/kg) was effective in increasing locomotor activity in both PERK^f/f^ DAT-Cre and control mice, as well as, in the drag (**Fig. 7d**), the rotarod (**Fig. 7e**) and the open field (**Fig. 7f**) test. Lastly, in one further test of DA involvement in the motor consequences of PERK deletion in DA neurons, we examined consequence of inhibition of the vesicular monoamine transporter 2 (VMAT2), which is also potentially relevant for pathophysiology in DA neurons, given that the ratio between DAT and VMAT2 is a vulnerability factor in DA neurons^37^. Interestingly, VMAT2 inhibition by low-dose reserpine (1 mg/kg) was sufficient to induce a variety of motor impairments in WT DAT-Cre mice (**Fig. 7d-f**), but affected only motor activity in the open field test (**Fig. 7f**) in PERK^f/f^ DAT-Cre mice. However, high-dose reserpine (1.5 mg/kg) administration resulted in altered motor behavior in both PERK^f/f^ DAT-Cre mice and controls (**Fig. 7d-f**) resulting in impaired stepping activity (**Fig. 7d**) rotarod performance (**Fig. 7e**) and distance moved (**Fig. 7f**). These data indicate that basic vesicular storage and release of dopamine is not impaired in PERK^f/f^ DAT-Cre mice. Taken together, these findings provide strong evidence that the changes in motor activity exhibited by the PERK^f/f^ DAT-Cre mice depends primarily on enhanced DA, and support the notion that similar patterns of DA release alteration may also underlie the changes in cognitive tests observed in PERK^f/f^ DAT-Cre mice.

**Figure 7.**
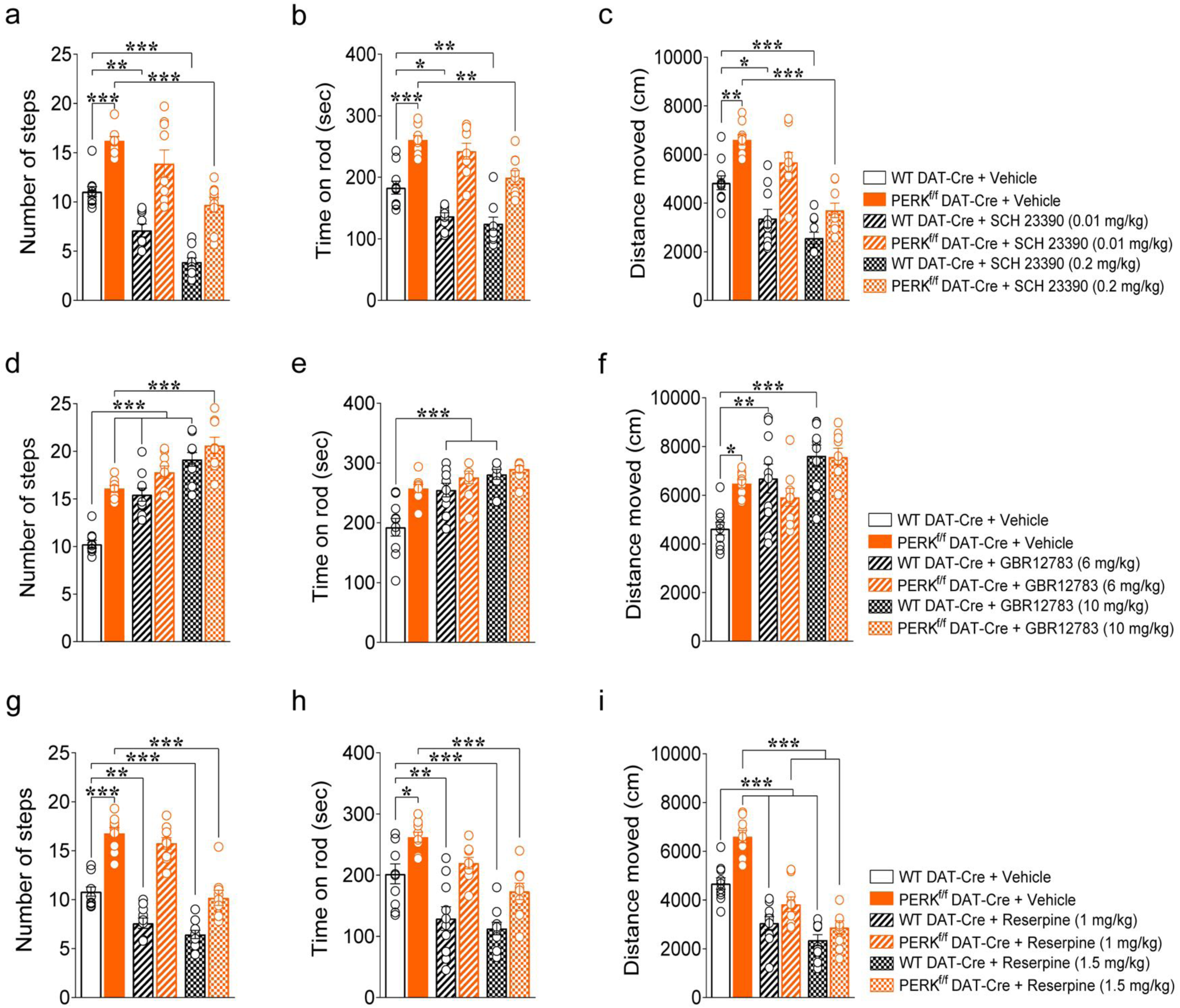
*In vivo* pharmacological targeting of DA machinery reveals the impact of PERK deletion in DA neurons in 3-month old mice. (**a-c**) Acute i.p. injection of the D1 antagonist (SCH 23390; 0.01 mg/kg) affects locomotor activity phenotype in WT DAT-Cre mice, but does not alter locomotor activity of PERK^f/f^ DAT-Cre mice. Conversely, high-dose SCH 23390 (0.2 mg/kg) impairs locomotor activity in both genotypes. (**a**) Summary plot of average number of steps during drag test (two-way ANOVA, genotype F*_(1, 51)_* = 93.91, P<0.0001, treatment F *_(2, 51)_* = 41.83, P<0.0001). (**b**) Summary plot of latency to fall from the rotating rod measured as average of two days (4 trials/day) test (two-way ANOVA, genotype F*_(1, 51)_* = 112.8, P<0.0001, treatment F*_(2, 51)_* = 18.19, P<0.0001). (**c**) Summary plot of spontaneous locomotor activity expressed as distance moved (cm) during the open field test (two-way ANOVA, genotype; F*_(1, 51)_*= 46.80, P<0.0001, treatment F*_(2, 51)_* = 34.66, P<0.0001). Mice were analyzed using two-way ANOVA followed by the Bonferroni’s test for multiple comparisons. *P < 0.05, **P < 0.01, ***P < 0.001. (**d-f**) Acute DAT blockade causes no changes in behavioral responses in PERK^f/f^ DAT-Cre mice, whereas increases locomotor activity in WT DAT-Cre mice at a lower dose. High-dose GBR12783 effect is consistent with greater DA uptake in PERK^f/f^ DAT-Cre mice. (**d**) Summary plot of average number of steps during drag test (two-way ANOVA, genotype x treatment, F*_(2, 51)_* = 6.63, P = 0.0028). (**e**) Summary plot of latency to fall from the rotating rod measured as average of two days (4 trials/day) test (two-way ANOVA, genotype x treatment, F*_(2, 51)_* = 4.12, P = 0.0221). (**f**) Summary plot of spontaneous locomotor activity expressed as distance moved (cm) during the open field test (two-way ANOVA, genotype x treatment; F*_(2, 51)_* = 5.59, P=0.0051). (**g-i**) VMAT2 blockade with i.p. administration of reserpine results in blunted behavioral responses in PERK^f/f^ DAT-Cre mice, in contrast to its impact on locomotor activity in control mice. (**g**) Summary plot of average number of steps during drag test (two-way ANOVA, genotype x treatment, F*_(2, 51)_* = 7.59, P = 0.0013). (**h**) Summary plot of latency to fall from the rotating rod measured as average of two days (4 trials/day) test (two-way ANOVA, genotype F*_(1, 51)_* = 39.58, P<0.0001; treatment F*_(2, 51)_* = 21.66, P<0.0001). (**i**) Summary plot of spontaneous locomotor activity expressed as distance moved (cm) during the open field test (two-way ANOVA, genotype x treatment; F*_(2, 51)_* = 4.118, P=0.0220). All data are shown as mean ± s.e.m. of n =10 (WT DAT-Cre) or 9 (PERK^f/f^ DAT-Cre) mice/treatment.

Previous studies have suggested that DA plays a crucial role in the induction and the modulation of striatal long-term depression (LTD),^38^ one of the two main forms of striatal synaptic plasticity^39, 40^ at corticostriatal synapses, which depends on the activation of DA receptors.^41^ In addition, reduction of evoked DA overflow in the striatum results in a failure to express LTD^42^. Therefore, we determined whether the altered *de novo* translation expressed in PERK-depleted SNc DA neurons underlying the changes in DA release, alters striatal LTD in cortico-striatal slices from 3 month-old PERK^f/f^ DAT-Cre and WT DAT-Cre mice. We recorded locally-evoked field excitatory postsynaptic potentials (fEPSPs) in the dorsolateral striatum, then delivered three trains of high-frequency stimulation (HFS) locally to induce LTD. We found that striatal LTD was enhanced in slices from PERK^f/f^ DAT-Cre compared to those from control mice (**Supplementary Fig. 5a,b**). DA plays a key role in the modulation of hippocampal synaptic plasticity and memory encoding, mostly through its binding to DA receptor^14^. Among the different sub-cortical inputs, VTA represents the major source of DA that the hippocampus receives^43^. Because our data from the NAc suggested changes in DA release in other VTA-innervated regions, such as the hippocampus and our behavioral results indicated that the disruption of PERK-eIF2α signaling in DA neurons alters hippocampus-dependent learning and memory in the PERK^f/f^ DAT-Cre mice (**Fig. 2**), we then examined late-phase long-term potentiation (L-LTP) in hippocampal slices from 3 month-old PERK^f/f^ DAT-Cre and control mice. We found that L-LTP, was enhanced in PERK^f/f^ DAT-Cre mice compared to WT DAT-Cre mice (**Supplementary Fig. 5c,dn**). Taken together, these findings suggest that PERK deletion in DA neurons significantly increases *de novo* protein synthesis in both striatonigral and mesocorticolimbic DA neuron populations, alters DA release and results in aberrant expression of both cortico-striatal LTD and hippocampal L-LTP, respectively.

### Selective disruption of PERK/eIF2α signaling in SNc DA neurons causes motor phenotypes similar to those exhibited by PERK^f/f^ DAT-Cre mice

DA neurons in the SNc and the VTA play pivotal roles in various brain functions, including the control of motor actions and higher cognitive functions such as learning and memory, motivation, decision-making, and reward processes^29^. These motor functions and cognitive abilities primarily involve two main DA pathways: the nigrostriatal and the mesocorticolimbic pathway, respectively. Thus, we determined whether disrupting PERK-eIF2α signaling in either the DA neurons of the nigrostriatal or the mesocorticolimbic pathway differentially impacted motor function and learning and memory at different ages.

We first investigated motor behavior following the selective deletion of PERK in SNc DA neurons. To conditionally delete PERK from DA neurons of the nigrostriatal pathway, we injected either AAV-TH-iCre ^44, 45^ or a control AAV expressing dsRED under TH promoter (AAV-Control) ^44^ into the SNc of 3-month old PERK^f/f^ mice (**Fig. 8a**). To verify the efficacy of PERK deletion, we examined co-expression of PERK and TH in PERK^f/f^ TH-dsRED AAV (control) and PERK^f/f^ TH-Cre AAV mice (**Fig. 8b-d**). In PERK^f/f^ TH-Cre AAV mice, PERK immunofluorescence was observed in ∼35% of TH^+^ cells (**Fig. 8c**). Notably, no difference was detected in the total number of TH+ cells between groups (**Fig. 8d**). We subjected PERK^f/f^ TH-Cre AAV mice and their controls to bar, drag, rotarod, NHC, and OF tasks (**Fig. 8**) 3 weeks after surgery. We found that 3-month old mice lacking PERK selectively in DA neurons of the nigrostriatal pathway exhibited a hyperactive motor phenotype expressed as reduced immobility time in the bar test (**Fig. 8e**) and higher stepping activity in the drag test (**Fig. 8f**). In addition, PERK^f/f^ TH-Cre AAV mice performed better than PERK^f/f^ TH-dsRED AAV mice in the rotarod task (**Fig. 8g**). Moreover, PERK^f/f^ TH-Cre AAV mice exhibited a significant increase in horizontal activity in the NHC task (**Fig. 8h**) when compared with the control mice. Although spontaneous locomotor activity in the OF task did not differ between PERK^f/f^ TH-Cre AAV mice and controls (**Fig. 8i**), PERK^f/f^ DAT-Cre mice did show an increase in vertical locomotor activity versus controls (**Fig. 8j**). We next determined whether deleting PERK in DA neurons of the SNc could affect learning and memory in 3-month old mice. We found a similar day-to-day decrease in escape latency between genotypes, during the acquisition of the hidden platform version of Morris water maze (**Supplementary Fig. 6a**). Also, no difference between groups was found in the time spent in the target quadrant (**Supplementary Fig. 6b**) and in the number of platform crossings (**Supplementary Fig. 6e**) in the probe test (**Supplementary Fig. 6a-b,e**), suggesting that hippocampus-dependent spatial memory deficits displayed by the PERK^f/f^ DAT-Cre mice (**Fig. 2**) is not due to disrupted PERK-eIF2α signaling in SNc DA neurons. Moreover, PERK^f/f^ TH-Cre-AAV mice exhibited similar performance compared to the PERK^f/f^ TH-dsRED-AAV control mice when tested in the novel object recognition (**Supplementary Fig. 6**) and in the LTM contextual threat memory (**Supplementary Fig. 6n**) tasks. Notably, PERK^f/f^ TH-Cre-AAV mice displayed a significant decrease in freezing when compared to the controls during the auditory cue (**Supplementary Fig. 6o**), denoting that selective deletion of PERK in SNc DA neurons plays a critical role in cognitive processes and affects amygdala-dependent associative memory in mice. Taken together, these results indicate that 3-month PERK^f/f^ DAT-Cre mice exhibit motor phenotypes due to altered function of the nigrostriatal pathway, and impaired associative threat memory that may be related to altered function of the central amygdala (CeA)-SNc-dorsal striatum (dStr) circuitry^46^.

**Figure 8.**
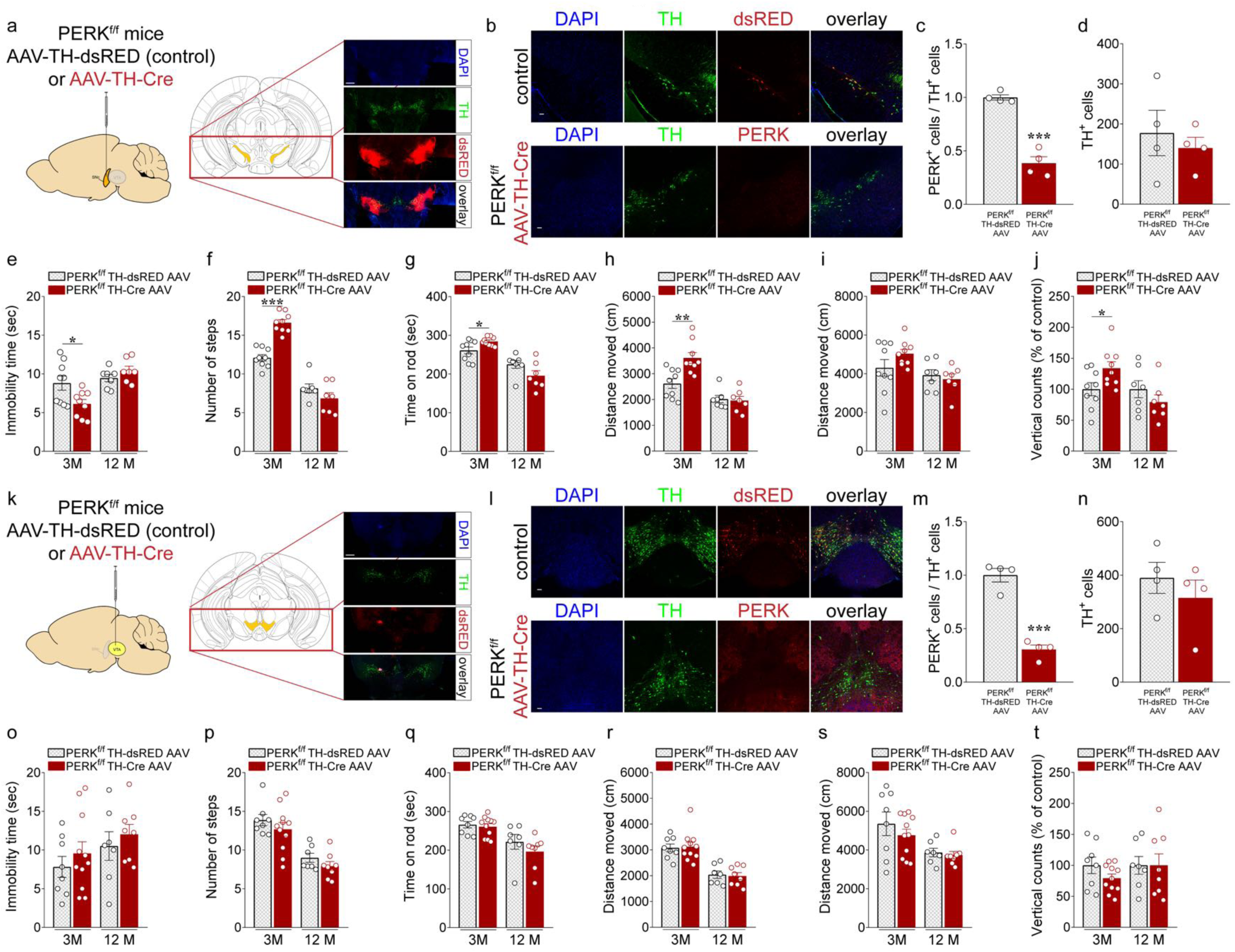
Mice lacking PERK in DA neurons of the SNc exhibit motor phenotypes similar to those displayed by the PERK^f/f^ DAT-Cre mice. (**a**) Schematic for experiments shown in **e-j**. Either AAV2/10 TH-dsRED (control) or AAV2/10 TH-iCre (AAV-TH Cre) was injected bilaterally into the SNc of PERK^f/f^ mice. Representative low magnification immunofluorescence images showing the injection sites (scale bar represents 500 µm). (**b**) Immunofluorescence detection of TH-IR (green) and either PERK or dsREd (red) in the SNc of PERK^f/f^ TH-Cre-AAV and PERK^f/f^ TH-dsRED AAV control mice (scale bars represent 50 µm). (**c**) Summary data showing the ratio of TH^+^ cells in the SNc that co-labeled for PERK in PERK^f/f^ TH-Cre-AAV *vs*. PERK^f/f^ TH-dsRED AAV control mice (*n* = 4 mice each, unpaired t test, t_(6)_ = 9.373, *P* < 0.001). (**d**) Summary data showing the overall number of TH^+^ cells in the SNc that co-labeled for PERK in PERK^f/f^ TH-Cre-AAV *vs*. PERK^f/f^ TH-dsRED AAV control mice (*n* = 4 mice each, unpaired t test, t_(6)_ = 0.599, *P* = 0.571). (**e-j**) PERK^f/f^ TH-Cre-AAV and PERK^f/f^ TH-dsRED AAV control mice were subjected to a set of tests including the bar (**e,o**), drag (**f,p**), rotarod (**g,q**), novel home cage (**h,r**), and open field (**i-j, s-t**) tests to investigate locomotor activity at 3 (3M) and 12 (12M) months of age. (**e**) Summary plot of immobility time (sec) during bar test in PERK^f/f^ TH-Cre-AAV *vs*. PERK^f/f^ TH-dsRED AAV control mice (unpaired *t* test, 3M: t_(16)_ = 2.500, *P* < 0.05; 12M: n.s.). (**f**) Summary plot of average number of steps during drag test (unpaired *t* test, 3M: t_(16)_ = 7.660, *P* < 0.001; 12M: n.s). (**g**) Summary plot of latency to fall from the rotating rod measured as average of two days (4 trials/day) test (unpaired *t* test, 3M: t_(16)_ = 2.439, *P* < 0.05; 12M: n.s). (**h**) Summary plot of the novelty-induced locomotor activity expressed as a distance moved (cm) in the first 10 minutes interval of a 60 minutes test during novel home cage test (unpaired *t* test, 3M: t_(16)_ = 3.586, *P* < 0.01; 12M: n.s). (**i-j**) Summary plot of (**i**) spontaneous locomotor activity expressed as distance moved (cm) and (**j**) vertical activity (number of counts) during the open field test (unpaired *t* test; **i**, n.s. at both 3M and 12M; **j**, 3M: t_(16)_ = 2.300, *P* < 0.05; 12M: n.s.). (**k**) Schematic for experiment shown in **o-t**. Either AAV2/10 TH-dsRED (control) or AAV2/10 TH-iCre (AAV-TH Cre) was injected into the VTA of PERK^f/f^ mice. Representative low magnification immunofluorescence images showing the injection sites (scale bar represents 500 µm). (**l**) Immunofluorescence detection of TH-IR (green) and PERK or dsRED (red) in the VTA of PERK^f/f^ TH-Cre-AAV and PERK^f/f^ TH-dsRED AAV control mice (scale bars represent 50 µm). (**m**) Summary data showing the ratio of TH^+^ cells in the VTA that co-labeled for PERK in PERK^f/f^ TH-Cre-AAV *vs*. PERK^f/f^ TH-dsRED AAV control mice (*n* = 4 mice each, unpaired t test, t_(6)_ = 9.120, *P* < 0.001). (**n**) Summary data showing the overall number of TH^+^ cells in the VTA that co-labeled for PERK in PERK^f/f^ TH-Cre-AAV *vs*. PERK^f/f^ TH-dsRED AAV control mice (*n* = 4 mice each, unpaired t test, t_(6)_ = 0.849, *P* = 0.429). (**o**) Summary plot of immobility time (sec) during bar test in PERK^f/f^ TH-Cre-AAV *vs*. PERK^f/f^ TH-dsRED AAV control mice (unpaired *t* test, n.s. at both 3M and 12M). (**p**) Summary plot of average number of steps during drag test (unpaired *t* test, n.s. at both 3M and 12M). (**q**) Summary plot of latency to fall from the rotating rod measured as average of two days (4 trials/day) test (unpaired *t* test, n.s. at both 3M and 12M). (**r**) Summary plot of the novelty-induced locomotor activity expressed as a distance moved (cm) in the first 10 minutes interval of a 60 minutes test during novel home cage test (unpaired *t* test, n.s. at both 3M and 12M). (**s-t**) Summary plot of (**s**) spontaneous locomotor activity expressed as distance moved (cm) and (**t**) vertical activity (number of counts) during the open field test (unpaired *t* test; n.s. at both 3M and 12M). All data are shown as mean ± s.e.m. of *n* = 9-7 PERK^f/f^ TH-Cre-AAV and *n* = 9-7 PERK^f/f^ TH-dsRED AAV mice injected in the SN at 3M and 12M, respectively; *n* = 11-8 PERK^f/f^ TH-Cre-AAV mice and *n* = 8-7 PERK^f/f^ TH-dsRED AAV mice injected in the VTA at 3M and 12M, respectively. \**P* < 0.05, \*\**P* < 0.01 and \*\*\**P* < 0.001 different from age-matched control.

We next determined whether PERK-eIF2α signaling disruption in the SNc DA neurons impacted motor and cognitive processes in 12-month old mice. We selectively ablated PERK in DA neurons of the nigrostriatal pathway in 12-month old PERK^f/f^ mice, using the same protocol described above. To our surprise, PERK deletion in SNc DA neurons did not affect motor function in these older mice (**Fig. 8**). PERK^f/f^ TH-Cre-AAV mice and their PERK^f/f^ TH-dsRED-AAV controls exhibited similar locomotor activity expressed as immobility time (**Fig. 8e**), stepping activity (**Fig. 8f**), and time on the rotarod (**Fig. 8g**) at 12 months of age. Moreover, no differences between groups were found in either horizontal (**Fig. 8h,i**) or vertical (**Fig. 8j**) locomotor activity in those mice. We then tested learning and memory in PERK^f/f^ TH-Cre-AAV mice and their PERK^f/f^ TH-dsRED-AAV controls at 12 months of age. We found that PERK deletion in SNc DA neurons in adult mouse brain affected neither hippocampus- and frontal cortex-dependent functions nor amygdala-dependent associative memory (**Supplementary Fig. 2**). Then, we examined striatal DA release in 12-month-old PERK^f/f^ TH-Cre AAV animals in which PERK omission was selectively restricted to either SNc or VTA DA neurons (*e.g.*, **Fig. 8**). Single-pulse evoked increases in [DA]_o_ were similar between control (PERK^f/f^ TH-dsRED AAV) and PERK^f/f^ TH-Cre AAV, in dStr, NAc core and shell, whether PERK was deleted selectively in SNc or VTA (**Supplementary Fig. 7**). Accordingly, no differences in the *V*_max_ for DA uptake were seen between genotypes for either injection site (data not shown). Similarly, midbrain DA levels were unaltered in PERKf/f TH-Cre AAV compared to the littermates at either age (**Supplementary Fig. 3d-f**). Collectively, these data suggest that the lack of PERK in DA neurons of SNc during early life alters control of motor function and importantly, that the age-dependent switch in motor performance in the PERK^f/f^ DAT-Cre mice requires PERK deletion during the aging process. Notably, alterations in the nigrostriatal pathway due to the selective deletion of PERK early in life might be detrimental for specific cognitive domains, such as associative threat memory.

### Selective disruption of PERK/eIF2α signaling in VTA DA neurons impairs learning and memory

Midbrain DA neurons of the VTA encode multiple signals that influence cognitive processes via diverse projections along mesolimbic and mesocortical pathways. Indeed, VTA projections to prefrontal cortex, hippocampus, amygdala and the ventral striatum are thought to regulate and contribute to various types of learning and memory^14, 31^. To conditionally delete PERK from DA neurons of the mesocorticolimbic pathway, we injected AAV-TH-iCre or the control AAV expressing dsRED under TH promoter (AAV-Control) into the VTA of 3-month old PERK^f/f^ mice (**Fig. 8k**). We verified the efficacy of PERK deletion by co-expression of PERK and TH in PERK^f/f^ TH-dsRED AAV (control) and PERK^f/f^ TH-Cre AAV mice (**Fig. 8l-m**). We observed PERK immunofluorescence in only ∼30% of TH^+^ cells of PERK^f/f^ TH-Cre AAV mice (**Fig. 8m**) and no difference was detected in the total number of TH^+^ cells between groups (**Fig. 8n**).

As expected, selective PERK deletion in DA neurons of the VTA did not alter locomotor activity in 3-month old mice. Indeed, PERK^f/f^ TH-Cre-AAV mice exhibited similar performance compared to the PERK^f/f^ TH-dsRED-AAV controls when examined in the bar (**Fig. 8o**), drag (**Fig. 8p**) and rotarod (**Fig. 8q**) tests. In addition, no difference was detected between genotypes in either horizontal (**Fig. 8r,s**) or vertical (**Fig. 8t**) locomotor activity. We proceeded to examine learning and memory in the 3-month PERK^f/f^ TH-Cre AAV and PERK^f/f^ TH-dsRED AAV mice. We found that PERK^f/f^ TH-Cre AAV mice displayed longer escape latencies (**Supplementary Fig. 6a**) compared to controls in the training phase of the MWM task. During the probe test, there was no difference between groups in the time spent in the target quadrant (**Supplementary Fig. 6b**), but the PERK^f/f^ TH-Cre AAV mice crossed the platform location significantly fewer times than PERK^f/f^ TH-dsRED AAV mice (**Supplementary Fig. 6e**). Moreover, we found that PERK^f/f^ TH-Cre AAV mice exhibited reduced preference for the novel object in both STM (**Supplementary Fig. 6h**) and LTM (**Supplementary Fig. 6i**) tests in the novel object recognition task. The selective deletion of PERK in the VTA DA neurons also recapitulated impaired associative memory deficits exhibited by the 3-month old PERK^f/f^ DAT-Cre mice in the associative threat memory tasks (**Supplementary Fig. 6**). PERK^f/f^ TH-Cre AAV mice displayed significantly decreased freezing time when exposed to the context (**Supplementary Fig. 6n**) and the auditory cue (**Supplementary Fig. 6o**) compared to their controls, consistent with the idea the disruption of PERK-eIF2α signaling in VTA DA neurons impacts several cognitive domains in a similar manner to 3-month old mice.

We proceeded to determine the effect of selective deletion of PERK via TH-Cre AAV injection in VTA DA neurons of the 12-month old PERK^f/f^ mice. PERK deletion in VTA DA neurons in these older mice also did not affect locomotor performance (**Fig. 8o-t**). Interestingly, however, the PERK^f/f^ TH-Cre AAV mice exhibited mild learning and memory impairments, mostly hippocampus-dependent, when compared to their controls (**Supplementary Fig. 6**). PERK^f/f^ TH-Cre AAV mice were trained to localize the hidden platform in the MWM tasks and they showed longer escape latencies (**Supplementary Fig. 6c**) compared to controls. Nevertheless, no difference between groups was found during the probe test (**Supplementary Fig. 6d,f**). The 12-month old PERK^f/f^ TH-Cre AAV mice showed altered STM (**Supplementary Fig. 6k**) and LTM (**Supplementary Fig. 6l**) in the novel object recognition task when compared with the PERK^f/f^ TH-dsRED AAV mice. Finally, we examined PERK^f/f^ TH-Cre AAV mice and their controls in the associative threat memory task. Both genotypes performed similarly during training, showing a similar freezing behavior (**Supplementary Fig. 6p**). However, PERK^f/f^ TH-Cre AAV mice exhibited a significant reduction in freezing time than controls when exposed to the context (**Supplementary Fig. 6q**), but no differences between groups were detected when the mice were exposed to the auditory cue (**Supplementary Fig. 6r**). These data suggest that hippocampus-but not amygdala-dependent memory processes were affected in the mice.

## Discussion

The upregulation of PERK-dependent UPR markers has emerged as a consistent feature of multiple neurodegenerative diseases over the last decade^9, 47^. There is evidence of dysregulated PERK-eIF2α signaling in *post-mortem* brain tissues of PD and AD patients^10^, as well as in preclinical models^48, 49^ of neurodegenerative disease. These studies marked a turning point in the recent research strategy on neurodegenerative disorders, bringing into focus the therapeutic value of UPR modulation across the spectrum of these diseases. Chronic activation of PERK signaling via eIF2α phosphorylation is thought to alter neuronal function by repressing global protein synthesis, particularly the synthesis and expression of a cluster of proteins important for the establishment of synapses and neuronal plasticity^18^. This type of PERK-eIF2α-dependent translation might account not only for synaptic failure, but also for and subsequent cognitive decline observed in most neurodegenerative disorders^24, 50, 51^. Thus, it is perhaps unsurprising that modulation of the PERK-eIF2α signaling has emerged as a promising therapeutic target for neurodegenerative disease. Much of the research focus regarding either pharmacological or genetic inhibition of PERK signaling to reduce p-eIF2α levels has been placed on models of pathology, suggesting a neuroprotective effect of restored eIF2α-dependent translation in mice ^18, 52–54^. However, there is other evidence from studies of preclinical models of neurodegenerative disorders that depict a complex scenario where, depending on the disease context, modulation of PERK-branch mediated translational control may result in contrasting and even opposite effects^19, 20, 55^. Thus, to fully understand the role of the UPR in pathology and normal brain function, dissecting the impact of modulating the PERK-eIF2α translational control pathway in a cell type-specific manner is necessary, especially if this pathway is to be harnessed as a therapeutic target to treat neurodegenerative diseases. In this study, we provide evidence that cell type-specific deletion of PERK in midbrain DA neurons results in multiple, age-dependent motor and cognitive phenotypes in mice. Our current study is the first demonstrating that sustained reduction of PERK-eIF2α signaling in DA neurons affects DA release and subsequently, DA signaling that impacts both nigrostriatal and mesocorticolimbic pathways.

Classically, eIF2α-mediated translational control in the brain has been studied in the context of synaptic plasticity as well as learning and memory processes^5,56^, with most of the studies focusing on molecular manipulation in excitatory neurons^17, 25, 26^. Our findings reveal a previously unrecognized role of PERK-eIF2α-mediated translation in DA neurons and provide molecular insights into the pathophysiology of motor and cognitive impairments when this type of translational control is disrupted. We found that reduced eIF2α phosphorylation in midbrain DA neurons leads to increased *de novo* translation, which results in DA dysfunction as well as multiple motor and cognitive impairments. Notably, there is an extensive body of literature describing the differential gene and protein expression between different DA neuron populations, which could underlie their selective vulnerability in the context of different neurodegenerative disorders^27, 57^. *De novo* protein synthesis was markedly increased in TH^+^ neurons of both the SNc and the VTA at 3 months of age, consistent with a clear dysfunction in both nigrostriatal and mesocorticolimbic pathways. Both pre- and postsynaptic protein synthesis is required for striatal LTD^39, 40^, which depends on the activation of DA receptors^40, 41^, as does locomotor behaviors, action selection, and associative learning^58^. The hyperactive motor phenotype exhibited by PERK^f/f^ DAT-Cre mice at 3-months of age is correlated with enhanced cortico-striatal LTD, likely *via* D2 receptor activation^59^ and elevated DA release in the dorsolateral striatum, as indicated by FSCV data (**Fig. 6**) and the pharmacological behavioral experiments targeting DA receptors and transporters (**Fig. 7**). Notably, preclinical models of PD, where degeneration of SNc DA neurons leads to striatal DA depletion, show impairment in indirect pathway mGluR-LTD^60^. The inhibition of indirect pathway LTD causes a shift of balance toward direct pathway LTP in the striatum, which ultimately inhibits movement^40^. Although DA binds to both D1 and D2 receptors, the relative activation of either subtype depends on the DA release levels and the respective affinities of the receptors for DA^29^. Either a hyper-dopaminergic state or a hypodopaminergic state can drive the system to become imbalanced, leading to unidirectional changes in plasticity that could underlie network pathology and symptoms^60^.

Despite no changes in DA content, PERK-deficient DA neurons showed increased DA release and uptake in the dStr and NAc of young PERK^f/f^ DAT-Cre mice that correlated with a marked hyperactive motor phenotype and required higher doses of SCH-23390 and GBR-12783 to block D1 receptors and DAT, respectively, than those sufficient to induce changes in behavior in WT DAT-Cre mice (**Fig. 7**). Conversely, a decrease in DA release and uptake in the dStr in older PERK^f/f^ DAT-Cre mice was associated with a marked decrease in locomotor ability, without affecting DA content (**Supplementary Fig. 3**). Our findings suggest that PERK not only is important for the regulation of DA release and DAT activity early in life, but sustained repression of PERK activity is detrimental for proper neuronal activity of SNc DA neurons during aging. Consistent with this notion, selectively deleting PERK from SNc DA neurons postnatally using a viral approach led to enhanced locomotor activity at 3 months of age, whereas no alteration in DA release and uptake, or in motor activity were detected when PERK was deleted in SNc DA neurons in 12-month old mice (**Supplementary Fig. 7**). Curiously, selective deletion of PERK in SNc DA neurons in young mice resulted in an amygdala-dependent memory impairment (**Supplementary Fig. 6**), which is consistent with the dense interconnections between the SNc and CeA identified as part of a CeA-SNc-CPu loop that underlies associative learning^46^.

It is well established that DA regulation of basal ganglia circuitry occurs mainly in striatum, the major input nucleus that also plays a central role in processing motivational, associative and sensorimotor information^61^. The increase in eIF2α-dependent translation via PERK deletion in VTA DA neurons is correlated with an increase DA release and uptake in the NAc of young mice (**Fig. 6**), whereas we detected an opposite outcome in the NAc when PERK was deleted in VTA DA neurons of 12-month old mice (**Supplementary Fig. 7**). DA plays a key role in the modulation of multiple forms of learning and memory by acting upon specific brain regions, including the hippocampus, amygdala, and prefrontal cortex, and altered DA signaling has been shown to impair the encoding and maintenance of memories^13–15^. Altered DA signaling resulting from PERK deletion in VTA DA neurons in both young and old mice may compromise the role of DA in the modulation of those target structures, ultimately resulting in impaired learning and memory. Indeed, DA is required for late-phase long-term potentiation (LTP) and spike timing-dependent plasticity in the hippocampus^62^, and DA receptor signaling regulates aversive memory retention^14^. Consistent with these observations, we found that hippocampal L-LTP is enhanced in young PERK^f/f^ DAT-Cre mice (**Supplementary Fig. 5**) and hippocampus-dependent contextual memory is altered in PERK^f/f^ DAT-Cre mice (**Fig. 2**), indicating that PERK activity in DA neurons is critical for encoding contextual information. Pharmacological activation of D1/D5 receptors, gates long-term changes in synaptic strength and facilitates induction and duration of LTP at CA1 and dentate gyrus synapses of the dorsal hippocampus in vivo^63^ and DAT re-uptake blockade, which increases DA availability, results in increased LTP magnitude in the CA1 region of rat hippocampus^64^. PERK deletion, specifically in DA VTA neurons in both young and old mice, results in spatial learning and memory impairments in the MWM task (**Supplementary Fig. 6**). In agreement with these findings, it has been shown that proper mesocorticolimbic DA neuronal function promotes hippocampal network dynamics associated with memory persistence^65^ and DA in the hippocampus has been shown to play a role in plasticity underlying spatial novelty^66^. The main DA input to the hippocampus arises from the VTA, which has been postulated to form a loop with the hippocampus that then regulates the activity of the VTA to control hippocampal activity through the release of DA^67^.

Moreover, deletion of PERK in excitatory neurons in the forebrain was shown previously to cause impaired cognitive function, especially in behavioral flexibility^25^. Finally, a reduction of eIF2α phosphorylation in either mice lacking the eIF2α kinase GCN2^50^ or in eIF2-S51A mutant mice^24^ results in a lowered threshold for the consolidation of long-term memory, impairments in long-term memory in standard learning and memory paradigms.

We also found that PERK^f/f^ DAT-Cre mice exhibit a preference for the familiar rather than the novel object in the novel object recognition test (**Fig. 2**), suggesting that impaired PERK-eIF2α signalling in VTA DA neurons results in prefrontal cortex information processing deficits. DA modulation in the prefrontal cortex is crucial for object recognition memory and the dysfunction of the dopaminergic system contributes to age-related cognitive decline in AD model mice^68^. Consistent with this notion, we showed that deleting PERK selectively in VTA DA neurons with a viral approach results in impaired spatial, associative, and discriminative memory in old mice (**Supplementary Fig. 6**). Our results suggest that PERK-eIF2α signaling is essential for proper function of mesecorticolimbic DA neurons as the mice age, and, therefore, it also plays a key role in the regulation of VTA DA neuron function in the adult DA system. Moreover, a sustained reduction of eIF2α phosphorylation, as is the case in aged mice, is detrimental for the integrity of DA signaling and either an increase or a reduction of DA outflow levels negatively affects DA neurons modulation of target structures. Notably, a recent study on cognitive deficits displayed by AD model mice pointed out the link between alteration in VTA DA neurons and deficits of hippocampus-dependent memory and synaptic plasticity^27^. It has been shown that a decrease in DA in the hippocampus and NAc shell of AD model mice due to VTA DA neuronal degeneration results in impaired synaptic plasticity, memory performance, and food reward processing, which suggests that altered VTA DA neuron function contributes to cognitive deficits in AD^27^.

UPR activation has been described as a “double-edged sword” because short-term activation plays a protective role whereas sustained activation results in synaptic failure, impaired synaptic plasticity, and ultimately, cellular death^7, 8^. Interestingly, multiple studies on neurodegenerative disease support the notion that prolonging, rather than inhibiting, PERK-eIF2α signaling results in neuroprotective effects. For example, in the A53T alpha-synuclein mutation model of PD, activation of the PERK branch mediates a pro-survival response^69^ and blocking eIF2α dephosphorylation in mutant SOD1 G93A mice, an amyotrophic lateral sclerosis (ALS) mouse model, prevents motor neuron degeneration and aggregation of mutant SOD1^70^. Thus, enhancing the PERK pathway by selectively inhibiting GADD34-mediated dephosphorylation of eIF2α in mutant model of SOD1 mice appears to significantly ameliorate the disease condition^71^. Finally, a recent study on multiple forms of dystonia, a brain disorder associated with involuntary movement and DA deficiency, demonstrated reduced eIF2α signaling in DYT1 dystonia patient-derived cells and that enhancing eIF2α signaling restored abnormal corticostriatal synaptic plasticity in a DYT1 mutant mouse model^55^.

In closing, our findings reveal the importance of PERK-eIF2α-mediated translational control in DA neurons and its role in normal motor function and in learning and memory. Moreover, given the general agreement that preventing translational repression caused by increased eIF2α phosphorylation might be beneficial for treatment of cognitive deficits associated with neurodegenerative disorders, our study sheds light on the consequences of long-term disruption of PERK in DA neurons for motor and cognitive function, suggesting that cell type-specific dissection of the role of PERK, as well as the UPR, in different neurodegenerative diseases is required. Finally, our study is not merely limited to the evaluation of the effect of PERK-eIF2α-mediated translational control repression, but uncovers an entirely new biological link between PERK and DA neuronal function that is involved in motor behavior and cognitive processes. Further investigation is needed to elucidate how disruption of PERK-eIF2α signaling affects general and gene-specific translation in DA neurons to alter DA release and uptake and ultimately, motor and cognitive behavior.

## Methods

### Animals

All mice were housed in groups of 3–4 animals per cage in the Transgenic Mouse Facility of New York University and maintained in accordance with the US National Institutes of Health Guide for Care and Use of Laboratory Animals. The facility was kept under regular lighting conditions (12 h light/dark cycle) with a regular feeding and cage-cleaning schedule. Mice were all maintained on a C57/BL6 genetic background and all genotypes were determined by polymerase chain reaction (PCR).

Transgenic mice obtained by selective ablation of PERK in midbrain DA neurons were generated by crossing mice harboring floxed PERK gene, *eIF2αk3* (PERK^f/f^; generated as previously described; Zhang et al. 2002) with heterozygous DAT-Cre recombinase mouse line (Jackson Laboratory, stock number: 006660)^21^ expressing *Cre* recombinase inserted upstream of the first coding ATG of the dopamine transporter gene (*Slc6a3*; DAT). The resulting heterozygous mice (PERK^f/+^ DAT-Cre) were crossed with PERK^f/+^ mice in order to obtain PERK DA conditional knockout (PERK^f/f^ DAT-Cre) mice and, the respective wild-type (WT DAT-Cre) littermates mice used as a control.

Knock-in eIF2α^(S51A/ S51A)^ mice expressing transgenic floxed wild-type *eIF2s1* gene were kindly provided by Dr. Randal J. Kaufman and generated as previously described^72^. The generation of eIF2α^(S51A/S51A)^ DAT-Cre mice, where the serine 51 residue has been mutated to alanine (S51A) selectively in DA neurons required two stages of breeding. First, eIF2α^(S51A/+)^ DAT-Cre mice were obtained by crossing eIF2α^(S51A/ S51A)^ mice with heterozygous DAT-Cre recombinase mouse line (Jackson Laboratory, stock number: 006660). Second, eIF2α^(S51A/+)^ DAT-Cre mice were crossed with eIF2α^(S51A/+)^ to generate conditional phospho-mutant eIF2α mice eIF2α^(S51A/S51A)^ DAT-Cre mice and, the respective wild-type (WT DAT-Cre) littermates mice used as a control. The resulting eIF2α^(S51A/S51A)^ DAT-Cre mice express both the floxed wild-type and the mutated eIF2α gene in all cells except for DA neurons (DAT+ neurons), where only the mutated eIF2α(S51A/S51A) form is expressed since the wild-type eIF2α gene is excised by *Cre*.

CAG^floxStop-tdTomato^(Ai14) conditional reporter line (B6; 129S6-*Gt(ROSA)26Sor^tm^*^14^*^(CAG-tdTomato)Hze^*; Jackson Laboratory, stock number: 007914)^23^ and PERK^f/+^ DAT-Cre mice were crossed to generate PERK^f/f^/CAG^floxStop-tdTomato^DAT-Cre mice and PERK^+/+^/CAG^floxStop-tdTomato^ DAT-Cre (named CAG^floxStop-tdTomato^ DAT-Cre) mice expressing tdTomato fluorescence following Cre-mediated recombination in DAT^+^ neurons. Mice generated from this cross were used exclusively to confirm the specificity of the Cre-recombinase system for the DAT^+^ neurons and the consequential deletion of PERK in dopaminergic neurons by immunofluorescence.

### AAVs infusion

AAV2/10-TH-iCre and AAV2/10-TH-dsRED adeno-associated viruses (∼10^12^ infectious units ml^-1^) were kindly provided by Dr. Caroline Bass and were generated as previously described^44^. Briefly, mice were anesthetized with a solution of ketamine hydrochloride (100 mg/kg, i.p.) and xylazine (10 mg/kg, i.p.), mounted onto a stereotaxic apparatus and viruses were infused bilaterally at the rate of 0.1 µl/min. Microinjection needles were left in place for an additional 5 min to allow for diffusion of viral particles. PERK^f/f^ mice were injected at either 2 or 11 months of age and allowed three weeks to recover after surgery. Injection coordinates targeting the SNc or the VTA were as follows (with reference to bregma): −3.1 AP, ± 1.2 ML, −4.3 DV (SNc) or −3.50 AP, ± 0.35 ML, −4.50 DV (VTA).

### Experimental design

For all behavioral and molecular experiments, mice of either sex were used. Mice subjected to locomotor activity analysis were tested at 3, 8 and 12 months whereas cognitive skills were tested in 3 and 12-month old mice. PERK^f/f^ mice (3 or 12 months of age) were subjected to intracranial injections of AAV-TH-Cre or AAV-dsRED in either the SNc or the VTA and tested for both motor and cognitive skills 3 weeks after AAV infusions. All mice were acclimated to the testing room 30 min prior to each behavioral experiment and all behavioral apparatuses were cleaned with 30% ethanol between each trial. The experimenter was blind to genotype. All behavioral tests were performed starting with the least aversive (locomotor activity) and ending with the most aversive (associative threat memory task) task.

### Mouse behavior: bar test

Originally developed to quantify morphine-induced catalepsy, this test measures the ability of the animal to respond to an externally imposed static posture. It can also be used to quantify akinesia (i.e. time to initiate a movement) also under conditions that are not characterized by increased muscle tone (i.e. rigidity) as in the cataleptic/catatonic state. Mice were gently placed on a table and forepaws were placed alternatively on blocks of increasing heights (1.5, 3 and 6 cm). The time (in seconds) that each paw spent on the block (i.e. the immobility time) was recorded (cut-off time of 20 s). Performance was expressed as total time spent on the different blocks. The test was performed in two consecutive days^73^.

### Mouse behavior: drag test

This test gives information regarding the time to initiate (akinesia) and execute (bradykinesia) a movement. It is a modification of the ‘wheelbarrow test’, and measures the ability of the animal to balance its body posture with the forelimbs in response to an externally imposed dynamic stimulus (backward dragging). Animals were gently lifted from the tail leaving the forepaws on the table, and then dragged backwards at a constant speed (about 20 cm/s) for a fixed distance (100 cm). The number of steps made by each paw was recorded. Five determinations were collected for each animal. The test was performed on two consecutive days^73^.

### Mouse behavior: rotarod test

The accelerating rotarod task (UGO BASILE, Biological Research Apparatus) was used to test balance and motor coordination. The rotarod test was performed by placing mice on a rotating drum (3 cm of diameter), and measuring the time that each mouse was able to achieve walking on the top of the rod. The time at which each animal fell from the drum, touching the plate at the base of the rod, was recorded automatically. If the mouse stopped walking, the session was considered ended at the third full turn of the drum without movement. The speed of the rotarod accelerated from 4 to 40 RPM over a 5 min (300 sec) period. Mice were given 4 consecutive trials with a maximum time of 300 sec and a minimum of 15 min inter-trial for two consecutive days. The fall latency (expressed in sec) obtained from each of 4 trials of the two days was used for statistical analysis^74^.

### Mouse behavior: novel home cage test (NHC)

The NHC test was used to assess the spontaneous horizontal motor activity as novelty-induced exploratory response. Mice were placed in a 35 x 22 x 22 cm experimental cage with the floor covered with bedding. Locomotor activity (expressed in cm) was recorded over a 60 min period by using a computerized video tracking system (Noldus, EthoVision XT 13)^75^. The parameter tested was the total distance traveled during the test and in each of the 6 intervals of 10 min.

### Mouse behavior: open field (OF) test

The OF test was used to measure the spontaneous general locomotor activity and anxiety-like behavior^76^. Mice were placed in the center of a clear Plexiglas open field (40 x 40 x 30 cm) for 15 min and a computer-operated optical system (Activity Monitor software for Open Field) monitored the movement of the mice as they explored in the open field. The parameters tested were: total distance traveled, vertical counts (expressed as % of control). The data were pooled according to genotype, and a mean value was determined for each group.

### Mouse behavior: novel object recognition task

The novel object recognition task was performed to analyze both STM and LTM^74^. Two familiar objects with same shape (2.5 x 2.5 x 2.5 cm) and color were used during habituation days. Objects with different shape (2.5 x 2.5 x 5 cm) and color from the familiar object, were used as novel objects. STM novel object was rectangular shaped and LTM novel object was round shaped. Mice were habituated to the arena and to the two familiar objects (FO) for 2 consecutive days and for 10 min each day. The following day, mice were exposed to the arena with the same two identical objects for the third time and the interaction time with both the objects was recorded (Noldus, EthoVision XT 13) during a 10 min period. After 1 hour, mice were tested for STM and they were exposed to the arena with one FO and the novel object (NO) replaced the non-preferred familiar object. Interaction time with both the objects was recorded for 5 min period. On day 4, mice were tested for LTM and the STM novel object was replaced with the LTM novel object; interaction time with both the objects was recorded for 5 min period and analyzed with the computer program. Interaction parameters were defined as contact with the object (noise-point detector; distance s; 2.54 cm). The Preference Index (PI) was calculated using the following equation:

*PI* = (*time interacting with* NO)/(*time interacting with* NO + *time interacting with* FO)

### Mouse behavior: Morris water maze

The Morris water maze test (MWM) was used to study spatial memory associated to a context. The maze consisted of a circular pool (52 cm in diameter) filled with water, and colored with white non-toxic paint to increase the water’s opacity. The pool was separated into four virtual quadrants. There were both proximal as well as distant visual cues in place. There were visual cues placed both on the pool and around the room to mark the position of the different quadrants and a hidden platform was placed in one quadrant. The diameter of the platform was 10.5 cm, which was 1 cm beneath the surface of the water. On days 1-5, mice were trained to locate the hidden platform. The training phase consisted of four (day 1) or 3 (day 2-4) trials per day with a cutoff of 60 sec and 15 min of inter-trials interval. On each day, the swim start positions for each trial were placed sequentially and counterclockwise in the quadrants, allowing mice to start from all the positions at the end of the training phase. The latency to find the platform was recorded. On day 5 mice were subjected to a 60 sec probe trial where the platform was removed from the pool. The parameters measured during the probe trial (also referred as test), were the number of platform crossings and time spent in each quadrant. One hour and a half after the probe trial, visible platform test was performed to ensure that none of the mice were impaired in their visual acuity. The platform was replaced in the same position as the training phase and a small flag was placed on the top of it to indicate the platform position, and the latency to find the visible platform was measured^74^. On day 6, the reversal learning task was performed. The hidden platform was placed in the opposite quadrant and the latency to find the new location of the platform was recorded. The reversal learning task continued for three days exactly as the initial training protocol (three trials each day; maximum duration of 60 sec for each trial)^77^. Escape latency, number of previous platform position crossings, time spent in each quadrant, and trajectories of the mice in both Morris water maze and water-based reversal learning task were recorded with a computerized video tracking system (Noldus, EthoVision XT 13).

### Mouse behavior: associative threat memory task

Contextual and cued threat conditioning (TC) was used to measure associative fear memory^78^. Mice were subjected to a neutral stimulus or context called “conditional stimulus” (CS), a 30 sec 80 dB tone; and an aversive stimulus called “unconditional stimulus” (US), a 2 sec 0.5 mA foot shock. Mice received two day of habituation (900 sec) to the training context, consisting of a white house light and metal conducting grid floor. On the training day mice were placed in the TC chamber (570 sec) and then given two sequences of CS-US, the first at 270 sec and the second at 420 sec. In this session mice associated the aversive experience (US) to the noise (CS). Freezing time was measured for intervals: from the start to the first CS-US (270 sec), during the first CS-US (30 sec), from the end of first CS-US to the second one (120 sec), during the second CS-US (30 sec) and from the end of second CS-US to the end (120 sec). On day 3, mice were probed first for context TC and then for cued TC, with one hour and a half interval. In context TC, mice were exposed to the training context as the habituation day (without foot shock and tone), and recorded for 300 sec; freezing time was measured every interval of 60 sec. In this session mice remembered the environment where they received the aversive stimulus. Cued TC was measured by placing mice in a novel context consisting of a red house light, a white Plexiglass floor and vanilla flavored bedding; cleaning solvents also was changed with 30% isopropanol. Mice were recorded for 720 sec and they were given three CS at 270 sec, 440 sec and 570 sec. Freezing time was measured in intervals: from 0 sec to 270 sec, during the first CS (30 sec), from the end of the first CS to the second CS (140 sec), during the second CS (30 sec), from the end of the second CS to the third CS (100 sec), during the third CS (30 sec), and finally from the end of the third CS to 720 sec (120 sec). Habituation, context TC and cued TC protocols, videos recording and freezing time analysis were performed with FreezeFrame^TM^ software (version 4.08, Actimetric Inc).

### Behavioral pharmacology

Three-month-old PERK^f/f^ DAT-Cre and WT DAT-Cre mice were acutely administered i.p. with the DAT inhibitor GBR-12783 dihydrochloride (Tocris Bioscience, cat no: 0421) at a doses of 6 or 10 mg/kg, the VMAT2 inhibitor reserpine (Tocris Bioscience, cat no: 2742) at a dose of 1 or 1.5 mg/kg, or the D1 receptor antagonist SCH 23390 hydrochloride (Tocris Bioscience, cat no: 0925) at a dose of 0.01 or 0.2 mg/kg, and motor activity was assessed using the drag, the rotarod, and the open field tests. Mice were tested 20 min (GBR-12783; SCH 23390) or 24 hr (reserpine) after drug administration. Experimenters were blind to the genotype and treatment.

### Fluorescent labeling of *de novo* protein synthesis in DA neurons

A fluorescent non-canonical amino acid tagging (FUNCAT) method was used to detect changes in *de novo* protein synthesis in DA neurons and it was performed as previously described^79^ with minor modifications. Briefly, 400 µm transverse slices were incubated with azido-homoalanine (AHA) at 32°C for 2.5 hours. At the end of the incubation slices were transferred into ice-cold 4% PFA for overnight fixation at 4°C. The following day, slices were mounted in 3% agarose and sliced using a vibratome (Leica VT1200S; Leica Microsystems; Bannockburn, IL) to a thickness of 30 µm. Free floating sections were collected in Tris-buffered saline (TBS), washed, blocked and permeabilized with 5% bovine serum albumin (BSA), 5% normal goat serum (NGS), 0.3% Triton-X-100 in TBS for 90 minutes (at RT). Slices were then washed with TBS and incubated with 500 µl of cyclo-addition reaction mix (Click-iT™ Cell Reaction Buffer Kit, Invitrogen, Ltd, Paisley, UK) for overnight cyclo-addition at 4°C with gentle rocking. Then, slices were rinsed in TBS, blocked in 1% NGS solution in TBS, and incubated with anti-tyrosine hydroxylase (TH) antibody (Millipore, MA, United States, cat no: MAB318). Slices were then rinsed in TBS and incubated with Alexa Fluor™488 goat anti-mouse secondary antibody (Invitrogen, Carlasbad, CA, USA). Finally, slices were rinsed with TBS and mounted using DAPI fluoromount-G™ (Electron Microscopy Sciences, Hatfield, PA, USA). AHA was detected using an Alexa Fluor™ 647 Alkyne, Triethylammonium Salt (Invitrogen, Carlasbad, CA, USA) and subsequent fluorescence imaging.

### Immunofluorescence and confocal microscopy

Mice were perfused intracardially with phosphate-buffered saline (PBS) followed by 4% paraformaldehyde. Brains were removed and stored in the same fixative o.n. at 4°C. The immunofluorescent staining was performed on 40-µm thick free-floating coronal midbrain slice containing VTA and SNc DA neurons that were prepared using a vibratome (Leica VT1200S; Leica Microsystems; Bannockburn, IL). For identification of DAT-specific deletion of PERK and consequential reduction of phosphorylated eIF2α (p-eIF2α) levels in VTA and SNc DA neurons, sections were blocked in 0.1% Triton-X-100, 5% NGS in TBS for 1h and incubated with primary antibodies o.n. at 4°C. The following primary antibodies were used: anti-tyrosine hydroxylase (TH) antibody (Millipore, MA, United States, cat no: MAB318) phospho-eIF2α (Ser51; Cell Signaling Technology, cat no: 33985), PERK (Cell Signaling Technology, cat no: 3192), anti-dsRED antibody (Thermofisher, cat no: 632496), anti-ATF4 (Santa Cruz, cat no: sc-390063). Alexa Fluor™ 488 or 647 secondary antibodies (Invitrogen, Carlasbad, CA, USA) were used. Sections were mounted using ProLong TM Gold Antifade mounting medium without DAPI (Invitrogen, Thermo Scientific, cat no: P36930). The sections were imaged using a Leica TCS SP8 confocal microscope (Leica, Germany) at 10X or 20X objective lens. All parameters (pinhole, contrast, gain and offset) were held constant for all sections from the same experiment. For analysis of AHA-Alexa-647 signal, 40 regions of interest (ROI) corresponding to soma size were selected from 20x TH-Alexa-488 images and were then measured in AHA-Alexa-647 images. Arbitrary fluorescent units (a.u.) were taken from Z-stack acquired images from the minimum value in the ROI to preserve the dynamic range as the increase in fluorescence between PERK^f/f^ DAT-Cre *vs*. WT DAT-Cre mice. For *de novo* protein synthesis in DA neurons detection the *n* refers to the number of mice per group (average of 40 somas per slice, 2 slices per mouse, from three independent experiments). All cell counting experiments were conducted blind to experimental group. All images were subsequently processed using ImageJ (NIH, USA).

### Protein synthesis assay

Proteins were labeled using a protocol adapted from the SUnSET method^80^. Briefly, 400 µm-thick brain slices containing SNc and VTA from 3-month old eIF2α^(S51A/S51A)^ DAT-Cre mice and control mice were prepared using a vibratome. Slices were allowed to recover in artificial cerebral spinal fluid (ACSF) at 32°C for 1 hour and subsequently treated with puromycin (P8833, Sigma-Aldrich, 5µg/mL) for 45 mins. Newly synthesized proteins were end-labeled with puromycin. SNc and VTA were micro-dissected from the brain slices and flash frozen on dry ice. Protein lysates were prepared for western blotting. Protein synthesis levels were determined by taking total lane density in the molecular weight range of 10 kDa to 250 kDa. Comparisons of protein synthesis levels between both genotypes were made by normalizing to the average WT DAT-Cre signal.

### Thapsigargin-induced ER stress assay

To induce ER stress, 400 µm-thick brain slices containing SNc and VTA from 3-month old mice were allowed to recover in ACSF at 32°C for 1 hr and subsequently treated with 1 µM Thapsigargin (Sigma-Aldrich, cat no: T9033) at 32°C for 2 hrs. ER stress was measured using either western blotting or immunofluorescent staining to detect PERK, p-eIF2α and ATF4 levels. SNc and VTA were punched out from the brain slices and processed by western blotting, whereas slices processed for immunofluorescence were embedded in 3% agarose solution and re-sliced at 40-µm thickness using a vibratome (Leica VT1200S; Leica Microsystems; Bannockburn, IL).

### Western blotting

SNc and VTA were separately micro-dissected from the brain slices and sonicated in ice-cold homogenization buffer (10 mM HEPES, 150 mM NaCl, 50 mM NaF, 1 mM EDTA, 1 mM EGTA, 10 mM Na4P2O7, 1% Triton X-100, 0.1% SDS and 10% glycerol) that was freshly supplemented with HALT protease and phosphatase inhibitor cocktail (Thermo Scientific, cat no: 78441, 1/10 total volume). Aliquots (2 µl) of the homogenate were used for protein determination with a BCA (bicinchoninic acid) assay kit (GE Healthcare). Samples were prepared with 5X sample buffer (0.25 M Tris-HCl pH6.8, 10% SDS, 0.05% bromophenol blue, 50% glycerol and 25% - β mercaptoethanol) and heat denatured at 95°C for 5 min. 40 µg protein per lane was run in pre-cast 4-12% Bis-Tris gels (Invitrogen) and subjected to SDS-PAGE followed by wet gel transfer to polyvinylidene difluoride (PVDF; Immobilon-Psq, Millipore Corporation, Billerica, USA) membranes. The membranes were probed overnight at 4°C using primary antibodies against p-eIF2α S51 (rabbit, Cell Signaling cat no: 9721, 1:500), anti-t-eIF2α (rabbit, Cell Signaling, cat no: 9722, 1:500), anti-puromycin (mouse, Millipore, cat no: MABE343, 1:1000). An antibody against β tubulin (mouse, Sigma, cat no: T8328, 1:1000) was used to estimate the total amount of protein. The membranes were probed with horseradish peroxidase-conjugated secondary IgG (1:7000, Promega) for 1 hr at room temperature. Signals from membranes were detected with ECL chemiluminescence (GE Healthcare Amersham™) using Alpha Imager 3.4 software and the FluorChem Protein Simple instrument. Exposures were set to obtain signals at the linear range and then normalized by total protein and quantified via densitometry using ImageJ software (NIH, USA).

### Electrophysiology

Striatal and hippocampal slice preparation and the recording of extracellular field excitatory postsynaptic potentials (fEPSPs) were performed as described previously^76^. Briefly, striatal and hippocampal slices from mice 3 to 4 months of age were isolated and transferred to recording chambers (preheated to 32 °C), where they were superfused with oxygenated artificial cerebrospinal fluid (ACSF). In all experiments, basal fEPSPs were stable for at least 20 min before the start of each experiment and all slices recovered in the recording chamber at least 1 hr before recordings began. Briefly, three trains of high-frequency stimulation (3 sec duration, 100 Hz frequency at 20 sec intervals) were used to induced LTD in striatal slices, while hippocampal L-LTP was induced with three 1 s 100-Hz high-frequency stimulation trains, with an intertrain interval of 60 sec^81^. After induction of either striatal LTD or hippocampal L-LTP, fEPSPs were collected for an additional 70 min and 140 min, respectively. Slope values of fEPSP were expressed as percent of the baseline average before LTD or L-LTP induction.

### Voltammetric monitoring of DA release using FSCV

Mice were deeply anesthetized with isoflurane, decapitated and brains were sliced with a Leica VT1200S vibratome (Leica Microsystems; Bannockburn, IL). Coronal forebrain slices (300 µm thickness) were cut in an ice cold HEPES-buffered artificial cerebrospinal fluid (aCSF) that contained (in mM): NaCl (120), NaHCO_3_ (20), glucose (10), HEPES acid (6.7), KCl (5), HEPES sodium salt (3.3), CaCl_2_ (2), and MgSO_4_ (2), bubbled with a 95% O_2_/5% CO_2_ gas mixture and recovered in this solution for 1 h at room temperature prior to recordings. Slices were transferred to the recording chamber, maintained at 32 °C and superfused (1.5 mL/min) with aCSF that contained (in mM): NaCl (124), KCl (3.7), NaHCO_3_ (26), CaCl_2_ (2.4), MgSO_4_ (1.3), KH_2_PO_4_ (1.3), glucose (10) equilibrated with 95% O_2_/5% CO_2_. Recordings were started after 30 min stabilization in the chamber. A Millar Voltammeter (available from Dr. Julian Millar, University of London, UK) was used for FSCV detection of evoked increased in [DA]_o_, as described previously^1,82^. Locally evoked DA release was detected using carbon fiber microelectrodes, constructed in house from 7-µm diameter carbon fibers (Goodfellow Corporation, Berwyn, PA, USA), with 30-70 µm exposed length. For FSCV, a triangular voltage ramp (from −0.7 to +1.3 V, then back to −0.7 V *vs*. Ag/AgCl) was applied every 100 ms at a scan rate of 800 V/s. Increases in [DA]_o_ in CPu and NAc core and shell were evoked using a concentric stimulating electrode positioned within 100 µm of the recording site. Single-pulse stimulation (400 µA, 0.1 ms duration) was used in CPu and NAc core, and a brief high-frequency pulse train (5 pulses at 100 Hz) was used in NAc shell to amplify evoked DA release. The currents resulting from DA oxidation were converted to DA concentration according to calibration with known concentrations of DA. To compare DA release in slices from control and PERK^f/f^ DAT-Cre or PERK^f/f^ TH-Cre mice, DA release was evoked in 4-5 sites in each striatal subregion in two slices per mouse. In all experiments after initial sampling under control conditions, a nAChR antagonist, DHβE (1 μM) was applied. A maximal drug effect was seen within 15 min, and multiple site sampling was repeated.

### DAT-mediated dopamine uptake analysis

The evaluation of changes in DAT activity was done using a MATLAB® script. *C*_peak_ values (peak concentration) and *V*_max_ values (maximum uptake velocity) for DAT-mediated dopamine uptake, derived from Michaelis-Menten analysis of single pulse evoked [DA]_o_ records, were determined as described previously^83^using a fixed *K*_m_ of 0.9 µM^84^ for each brain region and genotype. Data from the CPu and NAc core were used for this analysis because it was developed to extract uptake terms from [DA]_o_ transients evoked by single-pulse stimulation, which was used in these regions. It should be noted that all mice were heterozygous DAT-cre, which exhibit a small but significant impairment in DAT function^85^.

### HPLC analysis of striatal DA content

Quantification of DA content in experimental striatal slices was performed using HPLC with electrochemical detection, as described previously^82^. The dorsal and ventral striatum were dissected from each slice, with careful removal of overlying cortex and white matter. Samples (3-7 mg) were weighed, then frozen immediately on dry ice and stored at −80° C until processing. On the day of analysis, samples were diluted 1:10 in ice-cold mobile phase deoxygenated with argon, sonicated, centrifuged at 13,000 x g for 2 min and then the supernatant injected onto the HPLC.

### Statistical analysis

All data are presented as the mean ± SEM. Data was analyzed using GraphPad Prism 8. For behavioral experiments the experimenter was blinded to the genotype of the animals during behavioral testing. For two-group comparisons, statistical significance was determined by parametric and nonparametric two-tailed Student’s t tests. Multi-groups were analyzed using one-way ANOVA or two-way ANOVA. P values less than 0.05 were considered statistically significant. Extreme outliers were detected by applying Grubbs’ method with a = 0.05 to each experimental group and eliminated from further analysis (GraphPad software). Data distribution was assumed to be normal but this was not formally tested. For voltammetry experiments and C_peak_/V_max_ analysis *n* represents the number of recorded sites sampled from 3 to 5 mice per genotype. Statistically significant differences measured for WT *vs*. KO were assessed with two-tailed unpaired *t*-test with Welch’s correction for unequal variance in 12 month-oldmice during the C_peak_/V_max_ analysis. For this latter, values with *R*^2^ < 0.95 indicating goodness-of-fit were excluded from the data reported here. For the HPLC analysis, *n* is the number of samples analyzed for each region, with two to six samples per animal. Significance of differences was calculated using unpaired Student’s *t*-test. Statistical significance was considered to be *P* < 0.05.

### Data availability

The data that support the findings of this study are available from the corresponding author upon request.

## Acknowledgement

We thank Dr. C. E Bass (University of Buffalo) for providing the AAV2/10-TH-iCre and AAV2/10-TH-dsRED adeno-associated viruses and Dr. R. Kaufman (Sanford Burnham Prebys Medical Discovery Institute) for the Eif2(S51A) mouse line; We wish to acknowledge C. Farb for exceptional technical assistance and Dr P. Shrestha for critical advice and review of this manuscript. We thank all members of the Klann laboratory for critical feedback and discussions. The MATLAB script for Vmax analysis was written and provided by Dr. Charles Nicholson at NYU Grossman School of Medicine. This study was supported by National Institute of Health Grants (nos. NS034007 and NS047384) to E.K.

## Author contributions

F.L. carried out and analysed behavioral experiments, slice electrophysiology, and collected and analysed all in vivo and ex vivo data. M.M. carried out and analyzed FSCV experiments. S.A. carried out western blotting, and collected ex vivo data. P.L.I. and C.M. carried out behavioral experiments and collected in vivo and ex vivo data. J.C.P. performed DAT-mediated dopamine uptake analysis. M.E.R. coordinated, performed, and analyzed HPLC experiments. E.P. and N.R. carried out HPLC experiments. M.D. performed genotype detection. F.L., E.K., E.S. conceived the study. M.M. and M.E.R. participated in the design of the study. F.L. and E.K. designed and coordinated all the experiments and wrote the manuscript. All authors read and commented on the paper.

## Supplementary Figures

**Supplementary Figure 1.**
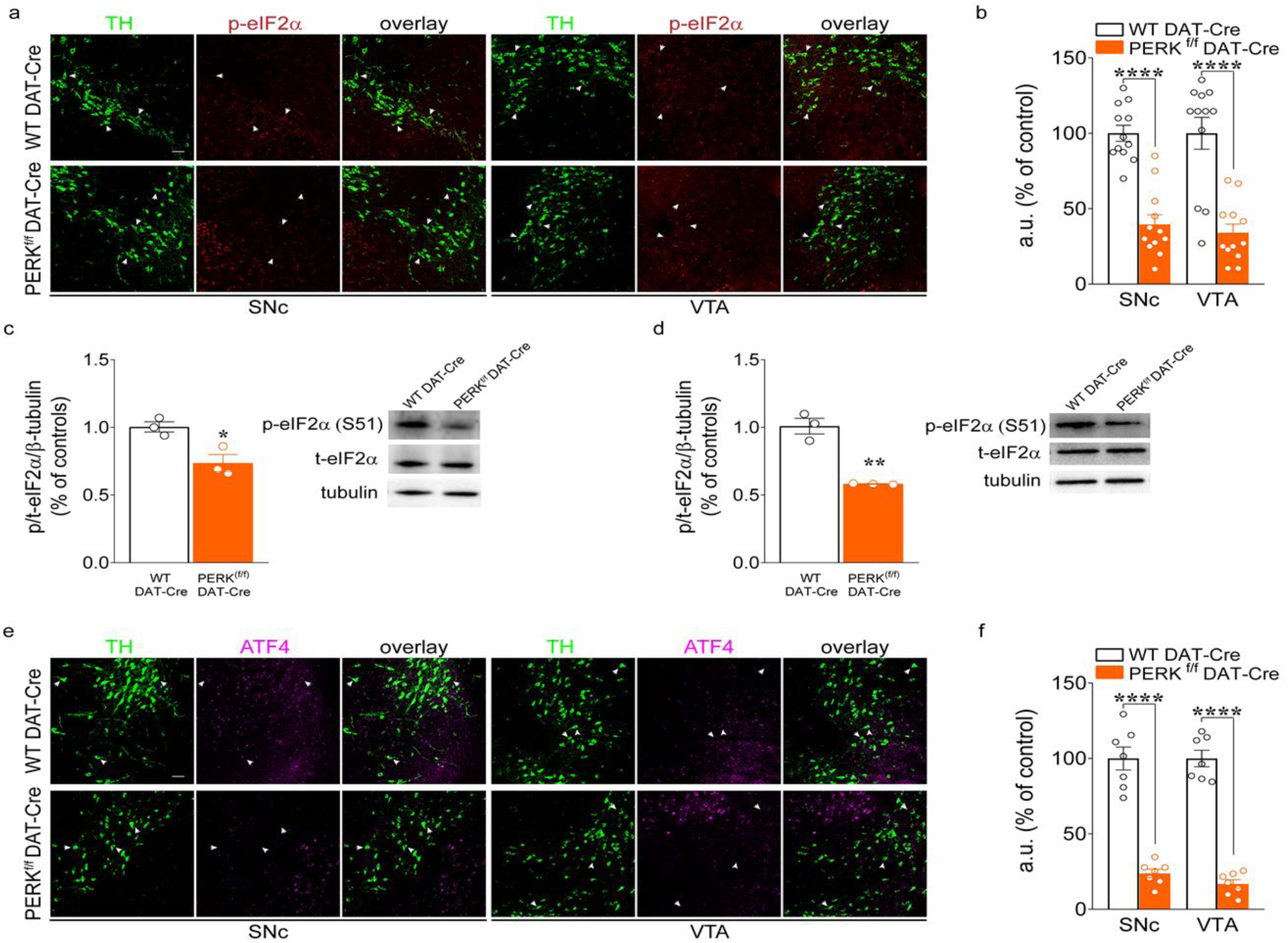
Cell-specific deletion of PERK in VTA and SNc DA neurons results in reduced UPR after thapsigargin-induced ER-stress. (**a**) Immunofluorescent detection of TH^+^ neuron (green) and phosphorylated eIF2α (red) in SNc and VTA DA neurons of PERK^f/f^ DAT-Cre and WT DAT-Cre mice, in thapsigargin-treated brain slices (scale bars represent 50 µm). Arrows indicate dopaminergic neurons (green) and p-eIF2α (red) co-stain. (**b**) Summary plot showing reduced p-eIF2α signal after ER-stress induction, expressed as fluorescent arbitrary units (a.u.; % of control) in TH+ neurons from SNc and VTA of PERK^f/f^ DAT-Cre vs. WT DAT-Cre mice. Cell soma intensity was measured in ImageJ. Statistical significance was determined by using Student’s *t* test (PERK^f/f^ DAT-Cre vs. WT DAT-Cre mice; unpaired t test; SNc, t*_(22)_* = 7.277, *P* < 0.0001; VTA, t*_(22)_* = 5.496, *P* < 0.0001). Data are shown as mean ± s.e.m. of n = 12 slices per group (average of n = 40 somas per slice, n = 2 slices per mouse, from three independent experiments) \*\*\*\**P* < 0.0001. (**c,d**) Representative western blots (right panel) and quantification of phosphorylation of eIF2α in SNc (**c**) and VTA (**d**) of PERK^f/f^ DAT-Cre and WT DAT-Cre mice. Summary plot showed a significant decrease in the phosphorylation of eIF2α in both SNc (**c**; unpaired t test, t*_(4)_* = 3.667, *P* = 0.023; n= 3 independent lysates from 3 mice per group) and VTA (**d**; unpaired t test, t*_(4)_* = 7.414, *P* = 0.002; n= 3 independent lysates from 3 mice per group) of PERK^f/f^ DAT-Cre mice compared with controls after treatment with thapsigargin. (**e**) Immunofluorescent detection of TH^+^ neuron (green) and ATF4 (magenta) in SNc and VTA DA neurons of PERK^f/f^ DAT-Cre and WT DAT-Cre mice, in thapsigargin-treated brain slices (scale bars represent 50 µm). Arrows indicate dopaminergic neurons (green) and ATF4 (magenta) co-stain. (**f**) Summary plot showing reduced ATF4 signal after ER-stress induction, expressed as fluorescent arbitrary units (a.u.; % of control) in TH+ neurons from SNc and VTA of PERK^f/f^ DAT-Cre *vs.* WT DAT-Cre mice. Cell soma intensity was measured in ImageJ. Statistical significance was determined by using Student’s *t* test (PERK^f/f^ DAT-Cre vs. WT DAT-Cre mice; unpaired t test; SNc, t*_(12)_* = 9.407, *P* < 0.0001; VTA, t*_(12)_* = 13.66, *P* < 0.0001). Data are shown as mean ± s.e.m. of n = 7 slices per group (average of n = 40 somas per slice, n = 2-3 slices per mouse, from three independent experiments) \*\*\*\**P* < 0.0001.

**Supplementary Figure 2.**
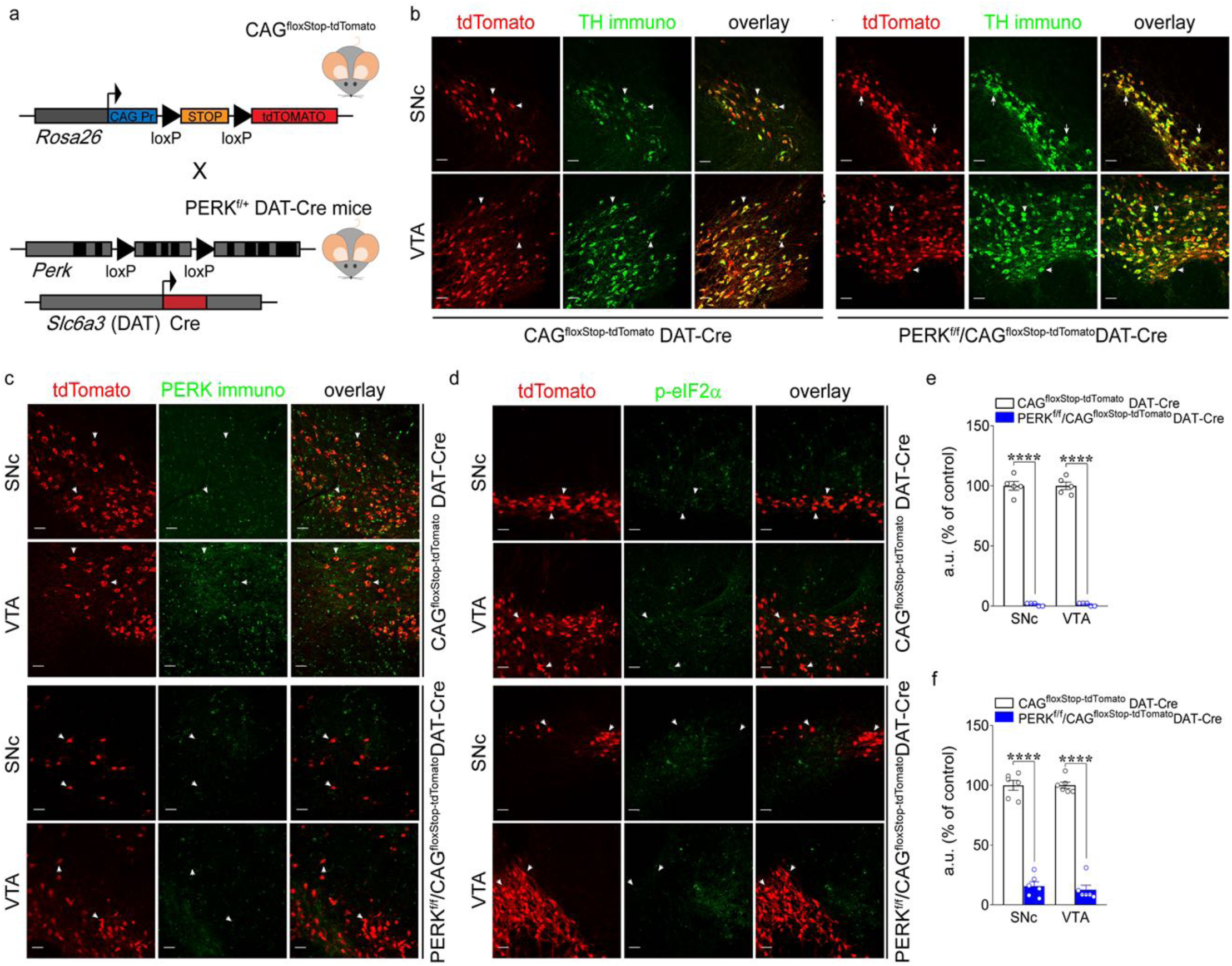
Gt(ROSA)26Sor^tm14(CAG-tdTomato)Hze^ Cre-dependent reporter mice crossed with PERK^f/+^ DAT-Cre mice confirm positive targeting of DAT^+^ neurons for the deletion of PERK and reduction of phosphorylated eIF2α in VTA and SNc DA neurons. (**a**) Schematic representation of DAT^+^ neuron-specific expression of tdTomato in DAT-Cre mice crossed with Gt(ROSA)26Sor^tm14(CAG-tdTomato)Hze^ Cre-dependent reporter mice. (**b**) Immunofluorescent detection of tdTomato (red) DAT^+^ neurons and TH-IR (green) in SNc and VTA DA neurons of PERK^f/f^/CAG^floxStop-tdTomato^DAT-Cre and WT/CAG^floxStop-tdTomato^DAT-Cre (control) mice, confirming specificity of the Cre-system (scale bars represent 50 µm). Arrows indicate dopaminergic neurons (green) and tdTomato (red) co-stain. (**c**) Immunofluorescent detection of tdTomato (red) DAT^+^ neurons and PERK (green) expression in SNc and VTA DA neurons of PERK^f/f^/CAG^floxStop-tdTomato^DAT-Cre and WT/CAG^floxStop-tdTomato^DAT-Cre mice, confirming positive targeting of DAT neurons for the deletion of PERK (scale bars represent 50 µm). Arrows indicate dopaminergic neurons (tdTomato; red) and PERK (green) co-stain. (**d**) Immunofluorescent detection of tdTomato (red) DAT^+^ neurons and phosphorylated eIF2α (green) expression in SNc and VTA DA neurons of PERK^f/f^/CAG^floxStop-tdTomato^DAT-Cre and WT/CAG^floxStop-tdTomato^DAT-Cre mice, confirming reduction of phosphorylated eIF2α following deletion of PERK in DA neurons (scale bars represent 50 µm). Arrows indicate dopaminergic neurons (tdTomato; red) and p-eIF2α (green) co-stain. (**e,f**) Summary plot showing reduced PERK (**e**) and p-eIF2α (**f**) signal expressed as fluorescent arbitrary units (a.u.; % of control) in tdTomato^+^ (dopaminergic) neurons from SNc and VTA of PERK^f/f^/CAG^floxStop-tdTomato^DAT-Cre *vs.* WT/CAG^floxStop-tdTomato^DAT-Cre mice. Cell soma intensity was measured in ImageJ. Statistical significance was determined by using Student’s *t* test (PERK^f/f^/CAG^floxStop-tdTomato^DAT-Cre *vs.* WT/CAG^floxStop-tdTomato^DAT-Cre mice; **e**, unpaired *t* test, SNc, t*_(8)_* = 27.04, *P* < 0.0001; VTA, t*_(8)_* = 31.14, *P* < 0.0001; **f**, unpaired *t* test, SNc, t*_(10)_* = 15.31, *P* < 0.0001; VTA, t*_(10)_* = 19.00, *P* < 0.0001) Data are shown as mean ± s.e.m. of n = 5-6 slices per group (average of n = 40 somas per slice, n = 2-3 slices per mouse, from three independent experiments) \*\*\*\**P* < 0.0001.

**Supplementary Figure 3.**
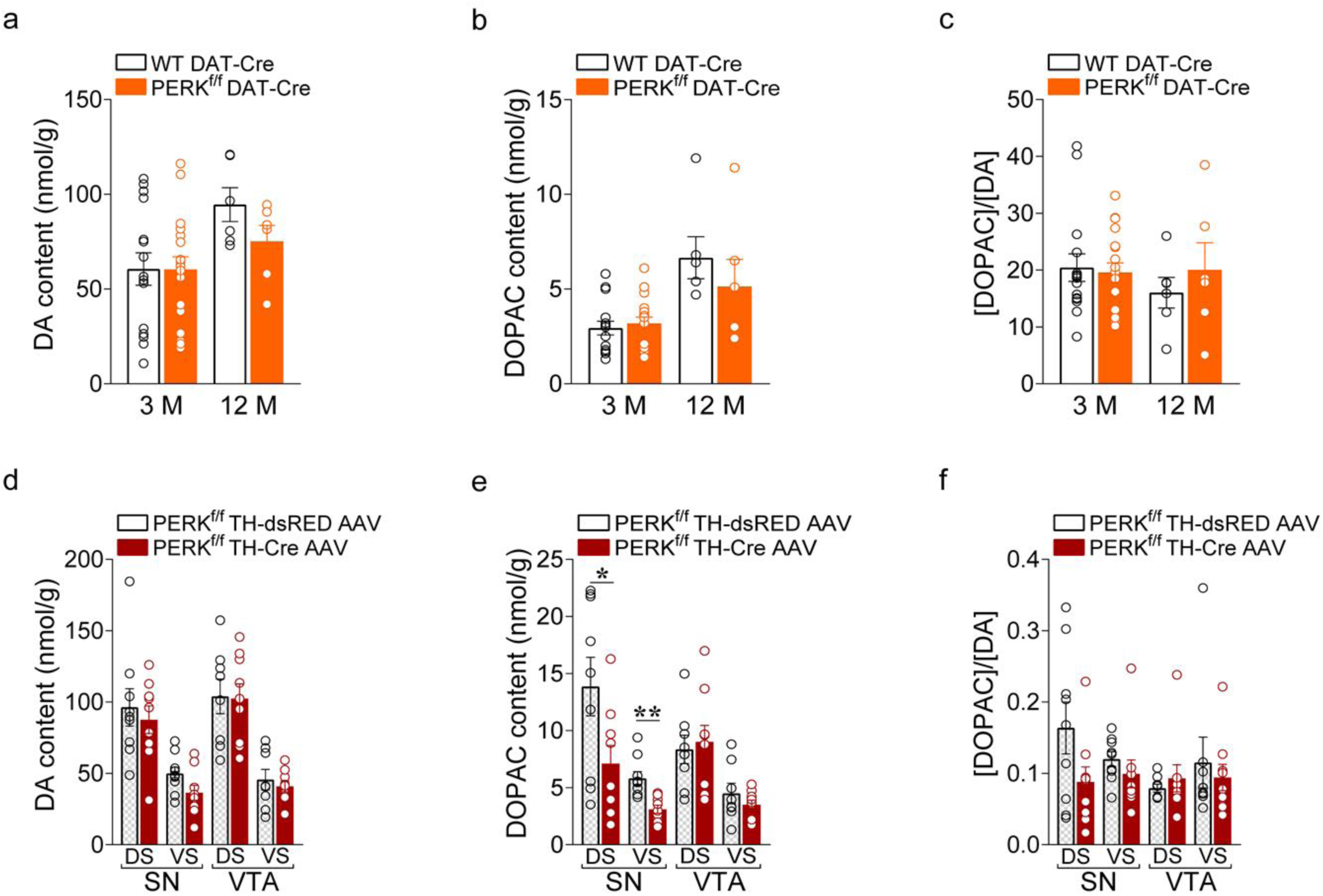
Deletion of PERK from DA neurons does not affect DA content in either 3- or 12-month old mice. Tissue content (nmoles/g tissue wet weight) of DA (**a,d**), its metabolite DOPAC (**b,e**), was performed using HPLC with electrochemical detection in experimental striatal slices. The analysis of the metabolite/DA ratio ([DOPAC/DA]; **c,f**) is also reported. (**a-b**) Summary plot of average DA (**a**) and DOPAC (**b**) content in both dorsal (DS) and ventral (VS) striatum of WT DAT-Cre *vs.* PERK^f/f^ DAT-Cre mice (unpaired *t*-test, n.s. in both 3 and 12-month old mice). (**c**) Summary plot of average [DOPAC/DA] ratio (unpaired *t*-test, n.s. in both 3 and 12-month old mice). (**d**) Summary plot of average DA content in either DS or VS of PERK^f/f^ TH-Cre-AAV *vs*. PERK^f/f^ TH-dsRED AAV mice injected in either the SN (unpaired t-test, n.s. in both DS and VS) or the VTA (unpaired t-test, n.s. in both DS and VS). (**e**) Summary plot of average DOPAC content in either in both DS or VS of PERK^f/f^ TH-Cre-AAV *vs*. PERK^f/f^ TH-dsRED AAV mice injected in either the SN (unpaired t-test, DS, t_(16)_ = 2.234, *P* < 0.05; VS, t_(16)_ = 3.936, *P* < 0.01) or the VTA (unpaired t-test, n.s. in both DS and VS) at 12 months of age. (**f**) Summary plot of average [DOPAC/DA] ratio in either DS or VS of PERK^f/f^ TH-Cre-AAV *vs*. PERK^f/f^ TH-dsRED AAV mice injected in either the SN (unpaired t-test, n.s. in both DS and VS) or the VTA (unpaired t-test, n.s. in both DS and VS). All data are shown as mean ± s.e.m. of *n* = 15-18 mice per genotype at 3 months of age and *n* = 6 mice per genotype at 12 months of age (a-c); or *n* = 8-9 TH-Cre AAV-injected 12-month PERK^f/f^ mice.

**Supplementary Figure 4.**
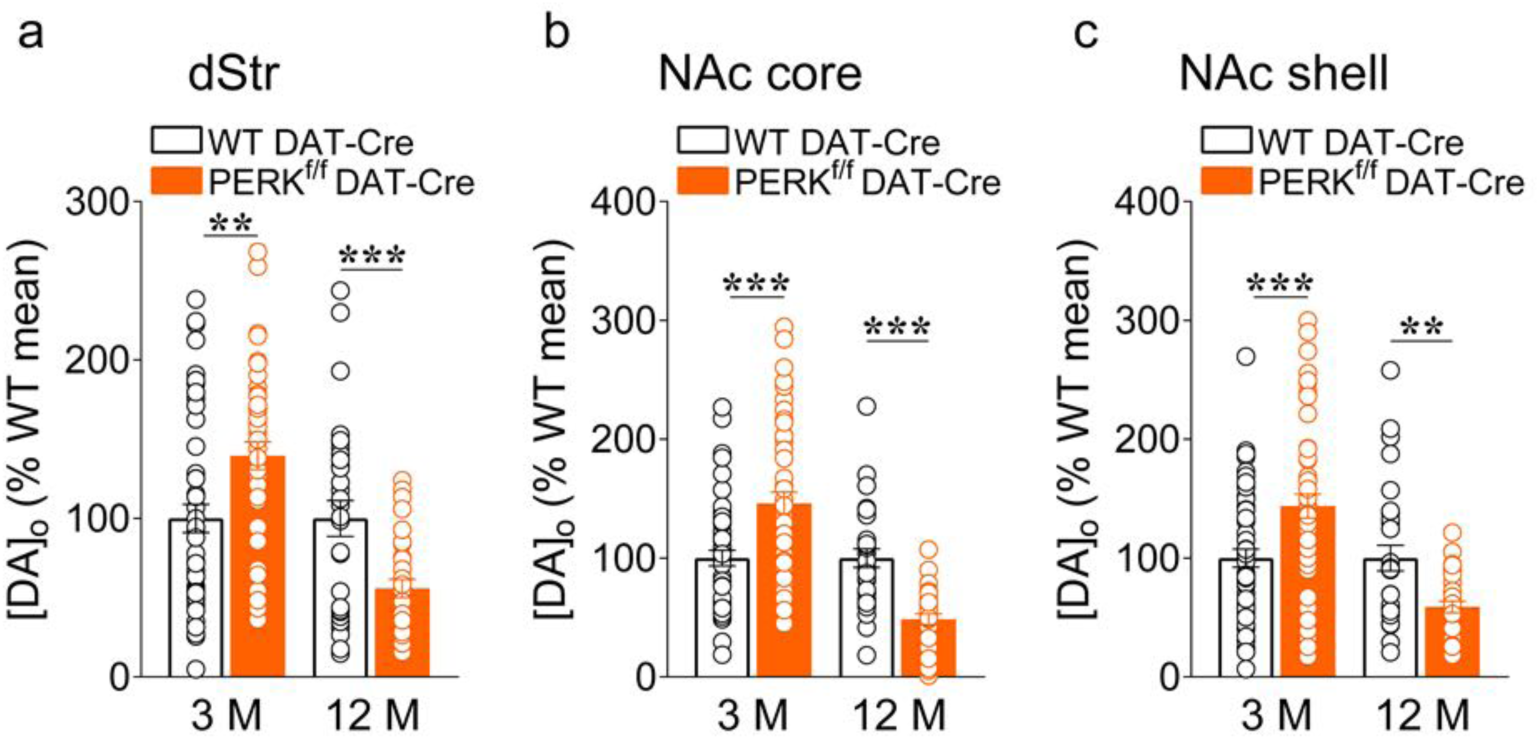
Effect of PERK deletion on DA release in DA neurons does not involve ACh and nAChRs. (**a-c**) Average single-pulse-evoked [DA]_o_ in dStr (**a**, unpaired *t-*test: t_(91)_ = 3.160, *P* < 0.01; t_(58)_ = 3.466, *P* = 0.001, respectively at 3 and 12 months), NAc core (**b**, unpaired *t-*test: t_(96)_ = 4.045, *P* < 0.001; t_(58)_ = 5.619, *P* < 0.001, respectively at 3 and 12 months), and NAc shell (**c,** unpaired *t-*test: t_(96)_ = 4.491, *P* < 0.001; t_(58)_ = 3.466, *P* = 0.001, respectively at 3 and 12 months) in the presence of dihydro-!3-erythroidine (Dh!3E; 1 µM), a selective antagonist for !32 subunit-containing (!32*) nAChRs that are enriched on DA axons. Data are are means ± s.e.m. normalized to 100% mean WT control for each region. \*\**P* < 0.01; \*\*\**P* < 0.001 for WT DAT-Cre *vs.* PERK^f/f^ DAT-Cre mice, where *n* denotes the number of recording sites in each region sampled from 3 to 5 mice per genotype.

**Supplementary Figure 5.**
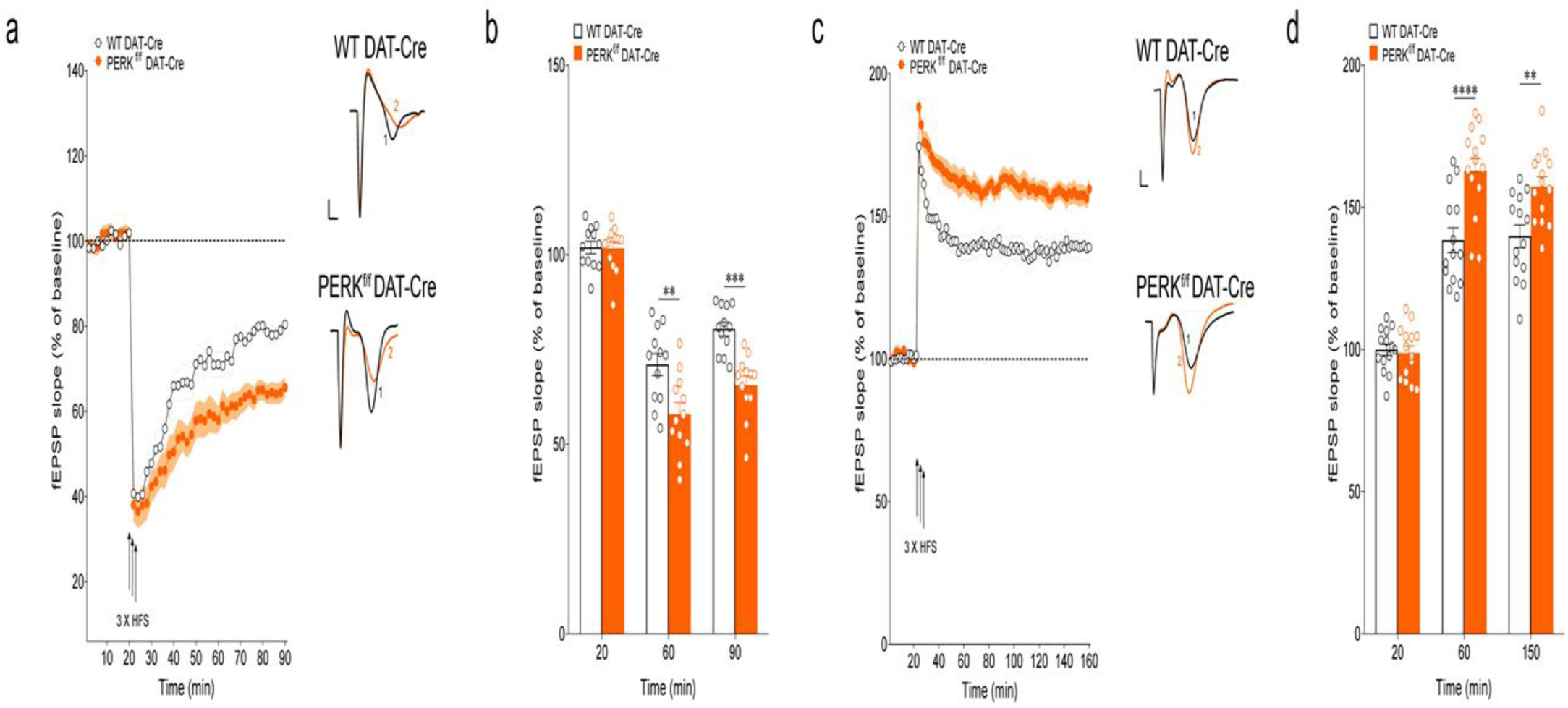
Deletion of PERK in DA neurons alters both striatal and hippocampal synaptic plasticity. (**a,b**) Striatal long-term depression (LTD) in 3-month old PERK^f/f^ DAT-Cre *vs*. WT DAT-Cre mice. (**a**) Plot showing normalized fEPSP mean slope (±s.e.m. displayed every 2 min) recorded from striatal slices. (**b**) Mean fEPSPs at baseline (20 min), at 60 (40 min after tetanus) and at 90 min (70 min after tetanus). LTD evoked by 3 trains of high frequency stimulation (HFS) was significantly enhanced in PERK^f/f^ DAT-Cre striatal slices at both 60 min and 90 min (two-way RM ANOVA, followed by Bonferroni’s multiple comparisons test, time x genotype, F*_(2,44)_* = 6.71, *P* = 0.003; time, F*_(2,44)_* = 161.60, *P* <0,0001; genotype F*_(1,22)_* = 19.85, *P* = 0,0002; n=12 slices from 7 mice/genotype). Data are shown as mean ± s.e.m. \*\**P* < 0.01 and \*\*\**P* < 0.001 PERK^f/f^ DAT-Cre *versus* WT DAT-Cre mice. (**c,d**) Hippocampal late-phase long-term potentiation (L-LTP) in 3-month old PERK^f/f^ DAT-Cre *vs*. WT DAT-Cre mice. (**c**) Plot showing normalized fEPSP mean slope (±s.e.m. displayed every 2 min) recorded from hippocampal slices. (**d**) Mean fEPSPs at baseline (20 min), at 60 (40 min after tetanus) and at 150 min (130 min after tetanus). L-LTP evoked by 3 trains of high frequency stimulation (HFS) was significantly increased in PERK^f/f^ DAT-Cre hippocampal slices at both 60 min and 150 min (two-way RM ANOVA, followed by Bonferroni’s multiple comparisons test, time x genotype, F*_(2,52)_* = 8.36, *P* = 0.0007; time, F*_(2,52)_* = 162, *P* <0,0001; genotype F*_(1,26)_* = 162, *P* = 0,0004; n=14 slices from 8 mice/genotype). Representative traces (**a,c** left panels) are superimposed fEPSPs (scale bars represent 1 mv/msec) recorded during baseline (1) and 60 min after HFS train (2). Arrows indicate delivery of HFS.

**Supplementary Figure 6.**
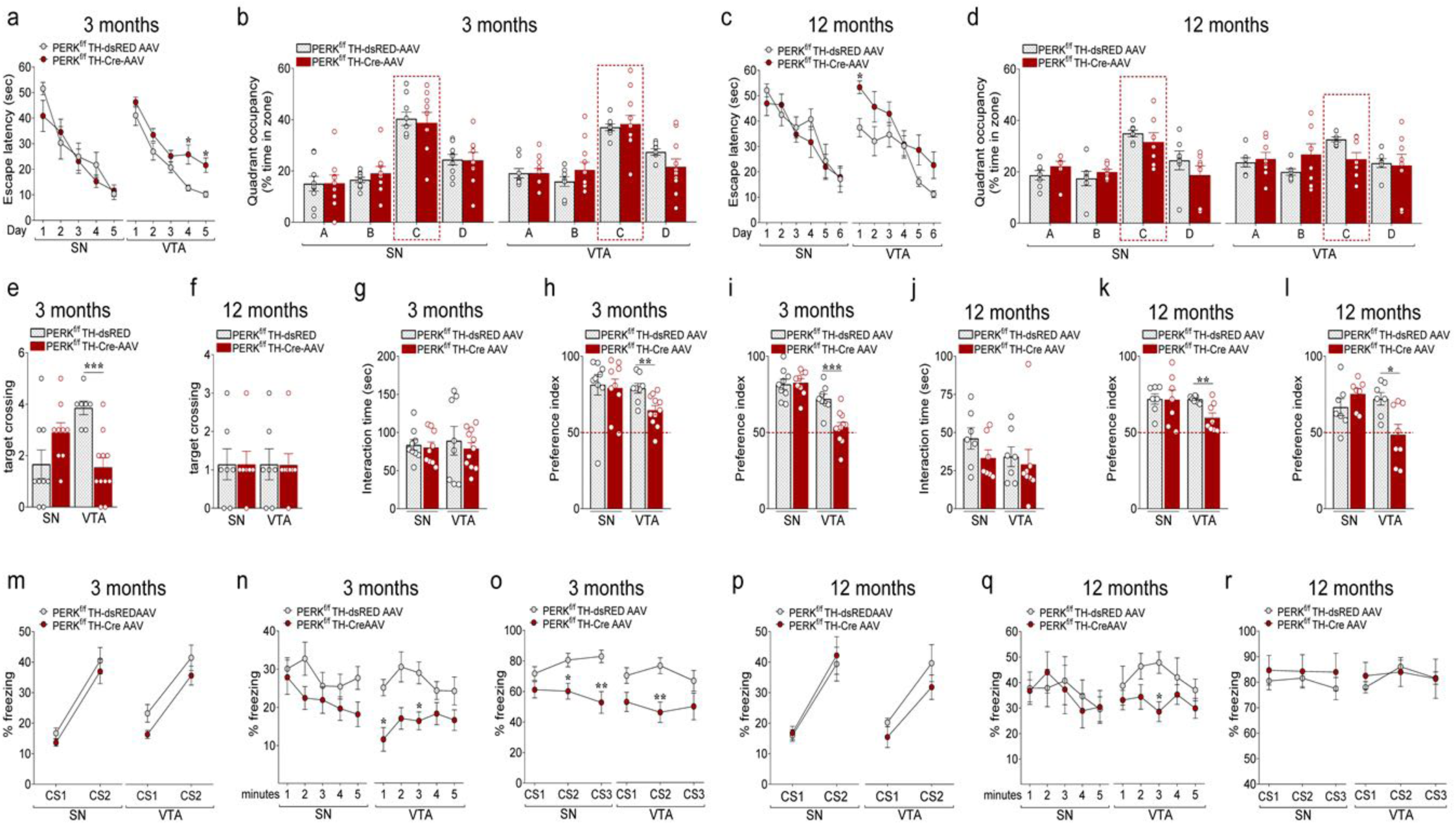
Mice lacking PERK in DA neurons of the VTA exhibit cognitive phenotypes similar to those displayed by the PERK^f/f^ DAT-Cre mice. (**a-f**) Summary plots of (**a,c**) average latency to find the hidden platform during a 5 (**a**) and 6-day (**c**) training protocol, (**b,d**) percentage of total time spent in each quadrant, and (**e-f**) average number of times spent crossing the location of the previously hidden platform during probe tests in 3-month old and 12-month old PERK^f/f^ TH-Cre-AAV *versus* PERK^f/f^ TH-dsRED AAV control mice injected in either SN or VTA in the MWM test (**a**, two-way RM ANOVA, followed by Bonferroni’s multiple comparisons test, SN: n.s; VTA: genoype, F*_(1, 17)_* = 15.22, *P* < 0.01; **b**, two-way RM ANOVA, followed by Bonferroni’s multiple comparisons test, quadrant x genotype, n.s., in both SN and VTA; **c**, two-way RM ANOVA, followed by Bonferroni’s multiple comparisons test, SN: n.s; VTA: genoype, F*_(1, 13)_* = 5.404, *P* = 0.037; **d**, two-way RM ANOVA, followed by Bonferroni’s multiple comparisons test, quadrant x genotype, n.s., in both SN and VTA; **e**, unpaired *t* test, SN: n.s.; VTA: t_(17)_ = 4.927, *P* < 0.001; **f**, unpaired *t* test, SN: n.s.; VTA: n.s.). (**g-l**) Summary plots of (**g,j**) interaction time with familiar objects, and (**e,f**) preference indices of mice toward a novel object introduced in the novel object recognition test in (**h-i**) 3-month old and (**k-l**) 12-month old PERK^f/f^ TH-Cre-AAV *versus* PERK^f/f^ TH-dsRED AAV control mice injected in either SN or VTA (unpaired t test; **g**, n.s. in both SN and VTA; **h**, SN: n.s., VTA: t_(17)_ = 3.173, *P* < 0.01; **i**, SN: n.s., VTA: t_(17)_ = 4.102, *P* < 0.001; **j**, n.s. in both SN and VTA; **k**, SN: n.s., VTA: t_(13)_ = 3.435, *P* < 0.01; **l**, SN: n.s., VTA: t_(13)_ = 2.866, *P* < 0.05). (**m-r**) Summary plots of average percentage of freezing during (**m,p**) training, (**n,q**) exposure to the context 24 hours after training, and (**o,r**) exposure to 3 CS presentations in a novel context in the associative threat memory test in (**m-o**) 3-month old and (**p-r**) 12-month old PERK^f/f^ TH-Cre-AAV *versus* PERK^f/f^ TH-dsRED AAV control mice injected in either SN or VTA (**m**, two-way RM ANOVA, followed by Bonferroni’s multiple comparisons test, n.s. in both SN and VTA; **n**, two-way RM ANOVA, followed by Bonferroni’s multiple comparisons test, genotype, SN: n.s., VTA: genotype, F*_(1, 17)_* = 10.97, *P* < 0.01; **o**, two-way RM ANOVA, followed by Bonferroni’s multiple comparisons test, SN: time x genotype, F*_(2, 32)_* = 5.837 *P* <0.01, VTA: time x genotype, F*_(2, 34)_* = 2.79 *P* =0.075; **p**, two-way RM ANOVA, followed by Bonferroni’s multiple comparisons test, n.s. in both SN and VTA; **q**, two-way RM ANOVA, followed by Bonferroni’s multiple comparisons test, genotype, SN: n.s., VTA: genotype, F*_(1, 13)_* = 11.22, *P* < 0.01; **r**, two-way RM ANOVA, followed by Bonferroni’s multiple comparisons test, n.s. in both SN and VTA). All data are shown as mean ± s.e.m. of *n* = 9 (3-month old) or 7 (12-month old) PERK^f/f^ TH-Cre-AAV and PERK^f/f^ TH-dsRED AAV mice injected in the SN; *n* = 11 (3-month old) or 8 (12-month old) PERK^f/f^ TH-Cre-AAV mice and *n* = 8 (3-month old) or 7 (12-month old) PERK^f/f^ TH-dsRED AAV mice injected in the VTA. \**P* < 0.05, \*\**P* < 0.01 and \*\*\**P* < 0.001 PERK^f/f^ TH-Cre-AAV *versus* PERK^f/f^ TH-dsRED AAV mice.

**Supplementary Figure 7.**
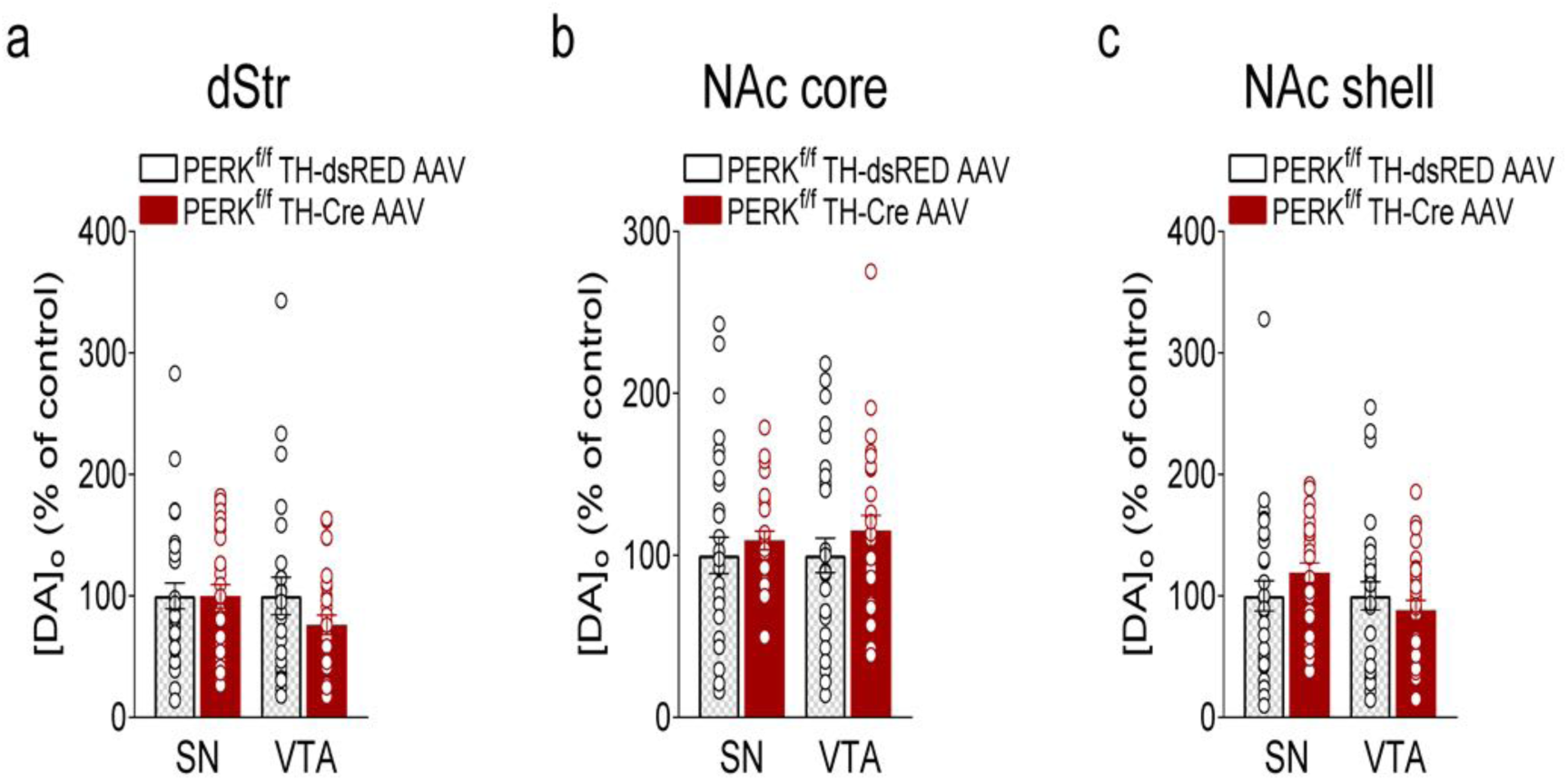
Selective deletion of PERK in DA neurons of either the VTA or the SN does not affect DA release and DAT activity in TH-Cre AAV-injected 12-month PERK^f/f^ mice. **(a-c)** Summary plot of evoked [DA]_o_ peak expressed as % of control in dStr (*a*, unpaired *t*-test, n.s. for either SN or VTA), NAc core (**b**, unpaired *t*-test, n.s. for either SN or VTA), and NAc shell (**c**, unpaired *t*-test, n.s. in either SN or VTA) in 12-month old mice. Data are means ± s.e.m. of *n* mice, where *n* denotes the number of recording sites sampled from 3 to 5 mice per genotype. Values with R^2^ < 0.95, indicating goodness-of-fit were excluded from the data reported here.

